# Developmental effect of RASopathy mutations on neuronal network activity on a chip

**DOI:** 10.1101/2023.09.20.558464

**Authors:** Eva-Maria Weiss, Debarpan Guhathakurta, Aneta Petrušková, Verena Hundrup, Martin Zenker, Anna Fejtová

## Abstract

RASopathies are a group of genetic disorders caused by mutations in genes encoding components and regulators of the RAS/MAPK signaling pathway, resulting in overactivation of signaling. RASopathy patients exhibit distinctive facial features, cardiopathies, growth and skeletal abnormalities, and varying degrees of developmental delay, neurocognitive impairment, intellectual disabilities, and attention deficits. At present, it is unclear how RASopathy mutations cause neurocognitive impairment and what their neuron-specific cellular and network phenotypes are. Such knowledge would be crucial for the development of specific therapies, which are still lacking for the treatment of the neurocognitive and behavioral symptoms in RASopathies. Here, we investigated the effect of RASopathy mutations on the establishment and functional maturation of neuronal networks. We isolated cortical neurons from RASopathy mouse models and cultured them for several weeks in multiwell plates with implanted multielectrode arrays. The arrays were used for longitudinal recordings of spontaneous activity in developing networks as well as for recordings of evoked responses in mature neurons. To facilitate the analysis of large and complex data sets resulting from long-term multielectrode recordings, we developed MATLAB-based tools for data processing, analysis and statistical evaluation. Longitudinal analysis of spontaneous network activity revealed a convergent developmental phenotype in neurons carrying the gain-of-function RASopathy mutations *Ptpn11*^D61Y^ and *Kras*^V14l^. The phenotype was more pronounced at early time points and faded out over time, suggesting compensatory effects during network maturation. Nevertheless, persistent differences in excitatory/inhibitory balance and network excitability were observed in mature networks. This study improves the understanding of the complex relationship between genetic mutations and clinical manifestations in RASopathies by adding insights into functional network processes as an additional piece of the puzzle.

## Introduction

RASopathies cover a group of genetic disorders associated with a characteristic pattern of anomalies, including distinctive facial features, cardiopathies, growth and skeletal abnormalities and various degrees of developmental delay [43]. They are typically caused by germline mutations in genes encoding components or regulators of the RAS/mitogen-activated protein kinase (MAPK) signaling pathway. Common is an overactivation in signaling, either by activating mutations in the pathway components or by loss of function in negative modulators [39, 43]. The RAS/MAPK signaling pathway has a central role in the cellular response to extracellular stimuli [27, 42]. In neurons, it is indispensable for proper differentiation, synapse formation and plasticity and is therefore critical for brain development, learning, memory and cognition [34]. Indeed, most RASopathies are associated with neurocognitive impairments and variable degrees of intellectual disability [7, 18, 38]. While patients with cardio-facio-cutaneous syndrome (CFC), Costello syndrome (CS) and SYNGAP1-related encephalopathy show moderate to severe intellectual disabilities and neurodevelopmental delay, epilepsy and autism spectrum disorder [1, 5, 30, 32, 36], patients with Noonan syndrome (NS) and neurofibromatosis type 1 are generally less severely affected. Common diagnoses are mild neurodevelopmental delay, impaired executive functioning, learning difficulties and attention deficit hyperactivity disorder symptomatology [2, 19, 21, 26, 29, 31, 37].

Interestingly, some of the neurocognitive features diminish with increasing age of the patients, while different features emerge. For example, children with NS have shown more extensive cognitive problems concomitant with learning difficulties and intelligence impairment compared to adults [33].

Animal studies have not yet addressed the developmental aspect systematically. While most studies demonstrated electrophysiological and behavioral phenotypes in mice carrying different RASopathy mutations, most of them did not assess the respective phenotypes along the animal lifespan [3, 10, 11, 23]. An exception is a recent study that used mice in which the RASopathy-related *Kras*^G12V^ mutation was expressed exclusively in neurons. In the hippocampi of these animals, upregulation of RAS/MAPK signaling was observed during the early phases of postnatal development but not in the adult state [28]. Interestingly, the normalization of RAS/MAPK pathway activity was concomitant with enhanced GABAergic synaptogenesis, indicating that maladaptive development of the neuronal network might underlie changes in RASopathy-induced neurocognitive effects observed during the lifespan in patients. Importantly, the switch in the cellular mechanism underlying network dysfunction might also have consequences for the selection of an effective treatment strategy. In line with that, Papale and colleagues were able to improve the cognitive phenotypes in *Kras*^G12V^ mice by treatment with RAS/MAPK inhibitors applied in juvenile animals. Strikingly, the same treatment was not effective if applied in older animals. Here, in line with increased GABAergic synaptogenesis, inhibition of GABAergic neurotransmission showed positive effects.

In this study, we follow up on the hypothesis that pathological activation of RAS/MAPK signaling potentially induces maladaptive processes leading to compensatory effects in developing neuronal networks. After all, neural circuits make up regulatory pathways for motion control, cognition, emotion, learning and memory in the brain. However, in vivo examination by real-time monitoring of neuronal activity and functional connectivity is restricted since deep tissue and components of neural circuits are difficult to access [6]. As an alternative opportunity to study functional connectivity and network parameters, we investigated neuronal cell cultures derived from RASopathy mouse models plated on multiwell MEA (mwMEA) plates in a longitudinal study. The disease models used in this study bear gain-of-function mutations in *Ptpn11*^D61Y^ [3] and in *Kras*^V14l^ [15, 17, 35]. To perform an extensive functional analysis of spontaneous network activity and electrically evoked activity, we used an in-house developed analysis tool written in MATLAB and routines for the application of principal component analysis (PCA) to reduce dimensionality in the data set for better data visualization and statistical evaluation. We observed significant effects of RASopathy mutation on network activity at early developmental stages that were compensated during network maturation. However, the mature RASopathy networks with apparently normal network activity still exhibited differences in excitatory/inhibitory balance and network excitability.

## Material and Methods

### Animals

For this study, conditional *Ptpn11*^D61Yfloxed/wt^ mice (B6.129S6-*Ptpn11*tm1Toa/Mmjax, Jackson Laboratories, RRID# MMRRC_032103-JAX) [8] and *Kras*^V14lfloxed/wt^ mice [17] were used. In these strains, a LOX-STOP-LOX cassette is cloned before the exon containing the RASopathy mutation. To generate a RASopathy mouse model, heterozygous mice bearing the mutation were crossed with mice homozygous for Emx1-cre alleles (B6.129S2-Emx1tm1(cre)Krj/J animals, Jackson Laboratories, RRID# IMSR_JAX:005628, genotyping according to provider’s recommendations). This strain expresses cre recombinase in forebrain excitatory neurons and glia from embryonic day 10.5 on. [14]. All mouse strains were bred on a C57BL/6 N background for more than 5 generations in our laboratory. Mice were kept at 22 ±2°C on a 12 h light-dark cycle with food and water ad libitum. Breeding of animals and experiments using animal material were performed in accordance with the local animal welfare officer (FAU: TS12/2016) and in accordance with the European Directive 2010/63/EU. Animal records were kept by Python-based Relational Animal Tracking Software (PYRAT-Scionics Computer Innovation GmbH, Dresden, Germany).

Animals were genotyped using published protocols. Specifically, the EMX1-cre genotyping protocol provided by the Jackson Laboratory was followed. To detect the *Ptpn11*^D61Y^ allele, polymerase chain reaction (PCR) using TGGAGCTGTTACCCACATCA and GCACAGTTCAGCGGGTACTT primers followed by a melting point analysis using the High Resolution Melting and Gene scanning application on the LightCycler 480 (Roche Diagnostics) was performed according to [3]. To determine the genotype in *Kras*^V14l^, PCR was performed using AGG GTA GGT GTT GGG ATA GC, CTC AGT CAT TTT CAG CAG GC, CTG CTC TTT ACT GAA GGC TC primers using S7 Fusion High-Fidelity DNA polymerase (Biozym, Cat#MD-S7-100) according to [17]. The protocol was generated to discriminate between 403 bp (wild type) and 621 bp (knock-in mutation) alleles. Primers were purchased from Eurofins, Ebersberg, Germany.

### Dissociated cell culture and plating on multiwell MEA plates or 18 mm coverslips

Chemical reagents are listed in Supplementary Material Table S 1. The day before dissection, wells of 48-well mwMEA plates (Cytoview Cat#MEA 48, M768-tMEA_48B, Axion Biosystems) and 18 mm Menzel glass coverslips (#6311342, VWR International, Radnor, USA) were coated with poly-L-lysine (0.5 mg/ml) followed by overnight incubation at 37°C. The next day, the wells and coverslips were washed three times with sterile double-deionized water and left in HBSS-/-, which was removed directly before cell seeding. Prior to seeding of cells on 18 mm coverslips, they were coated with 100 µl of Neurobasal^TM^-A media (NBA) supplemented with 0.2 mM Glutamax, 2% (v:v) B27, 0.1 M Antibiotic-Antimycotic (named NBA mix) and 10% fetal calf serum (FCS), as previously described [4], for better attachment of neurons.

Dissociated neuronal cultures were prepared from individual brains of newborn mice (P0-P1) as previously described in [4]. Briefly, *Ptpn11*^D61Yfloxed/wt^ (here termed *Ptpn11*^D61Y^) or *Kras*^V14lfloxed/wt^ (here termed *Kras*^V14l^) mice and their siblings (here termed control) were decapitated, the brains were removed, and the forebrains were separated and freed of meninges. Dissociation of cells took place chemically in Papain-protease-DNase mix (HBSS-/-, 0.01% Papain reconstituted in EBSS, 0.01% (w:v) DNase I, 0.1% (w:v) Dispase II) at 37°C for ten minutes followed by mechanical dissociation. This process was repeated 2 times. The cell suspension was passed through a 70 µm cell strainer. Cells were spun down and washed with NBA mix three times and then resuspended in NBA. Cells were counted and diluted to plate 80,000 cells in a volume of 50 µL onto the coated wells of the mwMEA plate. The cells were incubated at 37°C in a 5% CO_2_ atmosphere for 1 h and allowed to settle, followed by the addition of 250 µL of NBA mix for further culturing. Similarly, 200,000 cells in a volume of 100 µl were carefully seeded onto coverslips for 1 h immediately after removing the NBA mix containing FCS droplets. They were then transferred to 12-well plates containing 1 ml of NBA mix per well and maintained at 37°C in 5% CO_2_ until maturation. Genotyping was performed post hoc from tail biopsies obtained from euthanized pups.

### Immunocytochemistry and image acquisition

All antibodies used are listed in Supplementary Material Table S 1. Immunofluorescence staining of vesicular GABA transporter (VGAT) and vesicular glutamate transporter (VGLUT1) was performed as double staining in combination with integral synaptic vesicle protein synaptophysin. Neuronal cultures grown on coverslips for 21 days were fixed with 4% (w:v) paraformaldehyde in PBS for 4 min at room temperature (RT). Fixed neurons were then washed in PBS and blocked and permeabilized in tandem with PBS solution containing 10% (v:v) FCS, 0.1% (v:v) glycine and 0.3% (v:v) Triton X-100 for 45 min at RT (Supplementary Material Table S 1). Coverslips were incubated with primary antibodies at 4 degrees overnight and after washing with PBS with secondary antibodies at RT for 1 h. All antibodies were applied in PBS containing 3% (v:v) FCS. Coverslips were mounted on glass slides with Fluoroshield. All experimental conditions investigated in each experiment were processed in parallel with identical antibodies, solutions and other chemicals. Image acquisition was performed by an epifluorescence microscope (Nikon Eclipse Ti, Nikon Corporation) equipped with an iXon EM+ 885 EMCCD Andor camera (Andor Technology) using a 60X/NA 1.2 objective (Plan APO VC Nikon CFI, Nikon Corporation) and controlled by VisiView software (Visitron Systen GmbH).

### MEA recordings

The wells in 48-well MEA plates were equipped with 16 PEDOT electrodes arranged in a 4 x 4 grid with an electrode spacing of 340 µm, electrode diameter of 50 µm and recording area of 1.1 mm x 1.1 mm. Recordings were performed using a Maestro multiwell MEA recorder (Axion Biosystems) at 37°C in a 5% CO_2_:95% air atmosphere. Before recording, neuronal cultures were visually checked for vitality under a light microscope (Supplement Material Figure S 1A). Prior to each recording during the time series experiments, the MEA plate was kept in the MEA recorder for at least 5 min to allow the neuronal cultures to adjust to ambient conditions and to avoid bias by mechanical disturbances. To acquire data and for spike detection, we used Axion Integrated Studio (AxIS) software (2.4.2.13). The heatmap visualization in the software enabled a first control of neuronal activity online (Supplement Material Figure S 1B). Voltage potentials were recorded across all electrodes with a sampling rate of 12.5 kHz (Supplement Material Figure S 1C). Recordings were subdivided into 5-minute time units to avoid a large amount of data at once (.raw file). The raw signal was filtered by a Butterworth filter with a 200 Hz to 2.5 Hz bandpass filter. The filtered signal was detected for neuronal events when the signal exceeded a threshold denoted as spikes (cf. Ch. data analysis, Supplement Material Figure S 1D). These were stored in a spike list (.csv file) containing information about the time and amplitude of detected spikes chronologically.

#### Time series of spontaneous network activity

Multiple recordings of spontaneous activity in neuronal networks plated on the same mwMEA plate were longitudinally and noninvasively conducted in the time range 10 days in vitro (DIV) to 33 DIV. On a measuring day, spontaneous activity was recorded for 20 min in 5 min time units.

#### Disinhibition-induced stimulation experiments

The application of bicuculline (bic) to *Ptpn11*^D61Y^ neuronal cultures was performed on 33 DIV. The baseline was recorded for 20 min in 5 min time units, and then the plate was removed to add bic (volume: 10 µL, final concentration 10 µM, diluted in H_2_O from a 10 mM stock solution in DMSO). After treatment, the plate was placed back in the MEA recorder and kept there for an additional 10 min without recording, followed by 50 min post bic recordings in 5 min time units. (Supplementary Material Figure S 2). Values were normalized to the baseline (100%) and presented as percentages.

#### Electrical stimulation experiments

Electrical stimulations were applied on mature networks (three independent experiments on DIV29, DIV30 and DIV39). To provide electrical test stimulation (STIM) and tetanus stimulation (TET), individual electrodes from the MEA were triggered to deliver electrical pulses (Supplementary Material Figure S 3A). Protocols for STIM and TET were modified from [9] (Supplementary Material Figure 3C). The STIM protocol consists of a train of pulses (0.2 Hz). The pulses were biphasic from +750 mV to -750 mV lasting for 250 µs per phase. During STIM, six electrodes per array (stimulation electrodes e1-6, as shown in Supplementary Material Figure 3A) were triggered sequentially for 5 min, also denoted as sessions 1-6 (Supplementary Material Figure 3D). Prior to each STIM, which consequently lasted 30 min in total, spontaneous activity was recorded initially for 20 min (prior STIM1) and subsequently for 10 min (prior STIM2 and STIM3). (Supplementary Material Figure 3D). TET was delivered from one electrode and consisted of 20 bursts at 0.2 Hz containing 11 pulses at an intraburst frequency of 20 Hz (Supplementary Material Figure 3 C).

### Data analysis

#### Immunocytochemistry (ICC)

Identical camera and illumination settings were applied during image acquisition for coverslips of all experimental conditions imaged on the same day. Data were stored as 16-bit images and analyzed using the in-house MATLAB routine SynEval [16]. Herein, the pictures were segmented according to puncta labeled by immunofluorescence staining for synaptophysin (region of interest, ROI). Subsequently, within the ROI, puncta were identified by being positive for VGAT or VGLUT. In brief, the segmentation algorithm follows an iterative, water-shed-based approach, and the identification of puncta positive in costaining is based on the immunofluorescence intensity (IFI) threshold individually calculated for each picture and then applied to the averaged IFI value derived from ROI-encompassed pixels. Inter alia, the program returns the fraction and the IFI of positive puncta in costaining.

#### Processing MEA-based data

For further data analysis, customized software packages coded in MATLAB were used to process the spike lists (.csv) that were output by AxIS software. Each analysis was run in batch mode to save time and avoid human bias. The packages enable an analysis for each experimental setting, including spontaneous network activity in time series recordings plus network disinhibition experiments as well as electrically evoked activity. We extended the spontaneous network activity analysis for time series experiments with MATLAB programmed functions performing PCA commonly applied in machine learning to reduce dimensionality in data sets.

#### Spontaneous network activity

The software package for analyzing spontaneous network activity in longitudinal experiments is equipped with a graphical user interface (GUI) (Supplementary material Figure S 4) to enable the user-friendly input of experiment-related relevant data (cf. scheme in Supplementary Material Figure S 5A). In the first step, the user selects the spike list files (.csv) that had been output by Axion software after each measurement. The minimum spike rate to determine active electrodes (herein 0.1 Hz) and the minimum number of active electrodes (herein, unless otherwise stated: 10) can be specified in the GUI.

##### Spike Calculation

Spikes, which define neuronal firing events, are detected when the filtered voltage signal crosses a threshold in the continuous data stream. In this study, the threshold was adaptively set as a multiple of the median noise level (5.5 of standard deviation (SD)) on an electrode (Supplementary material Figure S 1C). The mean firing rate (MFR) is defined as the total number of spikes divided by recording time, MFR = n_spikes_/s. The weighted MFR is calculated as wMFR = (e_max_/ e_act_)·MFR. Herein, the maximum number of active electrodes (e_max_) was determined at the culture age of maturity (DIV 21-24). e_act_ refers to the currently active number of active electrodes. An electrode was defined as active when its spiking rate was at least 0.1 Hz, and then it was also described as a contributing channel. For disinhibition experiments, e_max_ was determined as the number of active electrodes during the baseline measurement prior to treatment. Analogously, wtMFR (mean firing rate weighted to total (t) number of electrodes) indicates the same calculation, but e_max_ refers to the total number of electrodes (herein 16 electrodes). The interpike interval is the mean time between spikes in s.

##### Burst detection and calculation

A burst was defined as neuronal activity that appears in high-frequency spiking followed by a time period of quiescence on a single electrode level. Herein, burst detection was performed along an interval interspike threshold detecting bursts as spiking events of no longer than 100 ms intervals between 5 consecutive spikes. The burst duration is calculated as the time from the first spike to the last spike within a burst. The mean bursting rate (MBR) is the total number of bursts within one well divided by recording time. The value is weighted analogously to wMFR (wMBR = (e_act_/ e_max_)·MBR).

##### Network analysis

The analysis of synchronized spiking in network bursts (NB) and correlated spiking measured as spike time tiling coefficient requires the transformation of the spike list to a time stamp matrix, a binary array of sample points indicating spikes as ones and sample points without spike as zeros. With a sampling rate of 12.5 kHz, one sample point correlates to 0.08 ms. The assessment of network-wide coordinated activity requires the detection of spike clusters across multiple electrodes.

Based on the time stamp matrix, a MATLAB-coded routine detected NBs across the array as spike clusters containing a minimum number of 50 spikes with a maximum of 100 ms interspike interval and with at least 5 contributing channels. Then, several NB-describing parameters, such as NB duration, number of NBs per well, and number of spikes per NB, were calculated. The NB duration was calculated as the time from the first spike to the last spike in an NB. To investigate changes in the network dynamics, the correlation of spiking between each pair of electrodes was analyzed by the spike time tiling coefficient (STTC) [12], which reveals the synchronization between pairs of electrodes by returning the unbiased correlation in firing. It is calculated as

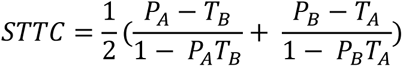

where A and B represent two spike trains. P_A_ (P_B_) is the proportion of spikes from A (B) that occurs within the time window ±Δt (100 ms) around any spike from B (A). T_A_ (T_B_) is the proportion of recording time that lies within ±Δt of any spike from A (B). The pairwise computation of STTC between electrodes resulted in a correlation matrix. For network description and illustration, we calculated the mean and skewness of STTC across all electrode pairs.

In the second part of the package (right side in GUI, compare Supplementary material Figure S 4c, d), functions are included that are capable of automatically grouping data according to a grouping file (.xlsx) that is provided by the user via the GUI.

#### Principal component analysis for time series recordings

For PCA, we developed functions in MATLAB that use the well-level data files obtained and saved after running the first part of the routine. Depending on the analysis, these functions created parameter arrays consisting of 19-dimensional parameter vectors (Supplementary Material Table S 2) to quantify cluster separation between the RASopathy model and wild type using principal components 1 and 2 (PC1/2 analysis). Furthermore, PCA was performed to create vectors of specific parameters describing the features spiking, bursting and network bursting (Supplementary Material Figure S 5B) following up the projection in PC1 with time. After PCA, MATLAB functions grouped well-level data automatically according to genotype or animal/preparation and time. For PC1/2 analysis, PCA score plots visualized genotype clusters in a PC1-PC2 two-dimensional coordinate system. To quantify the separation between the genotype clusters, we followed the suggested quantification method presented in [13]. In brief, we calculated the Mahalanobis distance [24] resulting in the distance between the centroids between groups considering groups’ data distribution. The calculations were performed by custom-written MATLAB functions that additionally calculated statistical significance with F-statistics.

#### Electrical evoked activity in stimulation experiments

For an automated analysis of data derived from MEA-based electrical stimulation of neuronal cultures, we developed a software package in MATLAB finally returning the network-wide evoked spiking rate upon STIM normalized to the spontaneous activity (evMFR_norm_) (data processing illustrated in Supplementary Material Figure S 6). Based on the initial measurement of spontaneous activity (SA1), preselection of valid wells was performed based on the condition of a minimum number of active electrodes per array (here: 8 electrodes) and with an electrode defined as active whenever its spiking rate exceeded a threshold (herein 0.1 Hz). To calculate evMFR_norm_, first, the evoked activity evMFR^’^was determined as the number of spikes within a time period of 1 s upon an electrical pulse divided by this time (Supplementary Material Figure S 3b), followed by averaging across all pulses on one electrode during one session. For each electrode, evMFR^’^ was then normalized to the spontaneous activity (MFR^′^_SA_) of the same electrode, resulting in evMFR_norm_. To evaluate the network-wide evoked activity, evMFR^′^_norm_ was averaged across all array electrodes and across all sessions within STIM, yielding evMFR_norm_. The software package returned a data array containing evMFR_norm_, as well as the corresponding coefficient of variance and the skewness for each well and STIM. Based on a user-provided Excel file, the package automatically grouped the wells according to genotype.

#### Data cleaning and selection of valid wells

To generate reliable results and to reduce the variability due to hardly predictable environment and physiological changes, criteria for the selection of valid wells were defined. First, wells were selected according to their activity as measured by the minimum number of active electrodes (if not otherwise stated: 10) determined by the minimum spiking rate (herein > 0.1 Hz). For the time series experiments, this selection was performed based on measurements taken at the time of network maturity, which we denote here as the reference time point (DIV 21-24). For disinhibition-induced stimulation experiments, the baseline measurement at the fourth time point was taken as a reference. Wells that did not fulfil the criteria were discarded from the analysis. We implied further quality criteria and discarded wells once they lost their integrity beyond the reference time point. Loss of integrity was measured as a considerable drop in the number of active electrodes (no. of contributing channels < 80% of no. of contributing channels at reference time point) or as a clear decrease in neuronal activity (wMFR < 70% wMFR_ref_).

For stimulation experiments that were more prone to interference due to a measurement duration above 2 h, additional exclusion criteria were applied to ensure that only stable cultures were included in the analyses. First, cultures had to exhibit stability in their spontaneous activity prior to (SA1) and after (SA2) STIM1 (Supplementary Material Figure S 3d). For each well, the MFR and SD of the firing rate were computed, and significant changes between SA1 and SA2 were identified by Welch’s t test to discard wells that were significantly altered. Second, of particular importance for the evaluation of the effect of tetanic stimulation on the network evoked activity, we discarded cultures that were pathologically affected by STIM sessions. To evaluate this, we regarded the relation between *evMFR*_*norm*_ upon STIM2 and upon STIM1 and checked for outliers (Supplementary Material Table S 4).

### Statistics

For statistical calculations, we used GraphPad Prism (version 8.3.0) Software and MS Excel. The distribution of all data sets was tested by normality and lognormality tests (Supplementary Material Table S 4 and Figure S 7).

To test for significant differences within time series experiments, subsequent PCA and disinhibition-induced stimulation experiments, we performed a mixed-effects analysis with GraphPad Prism. For time series experiments, we assume an increased risk of type two error caused by a systematic effect by time on the whole population potentially superimposing the effect of genotype and therefore performed either an unpaired t test for normally distributed data or Mann‒Whitney U tests for nonnormally distributed data without correction for multiple comparisons. The effect size is indicated as r^2^ for unpaired Student’s t test [22] or r for the Mann‒Whitney U test according to Wendt [41] (Figure 1+2). Standard benchmarks for r fit well with our set of data: <.3 small effect,.3-.5 moderate effect, and >.5 strong effect (r^2^: <.09 small effect,.09-.25 moderate effect, and >.25 strong effect, respectively). A p value <.05 was considered significant.

**Fig. 1.**
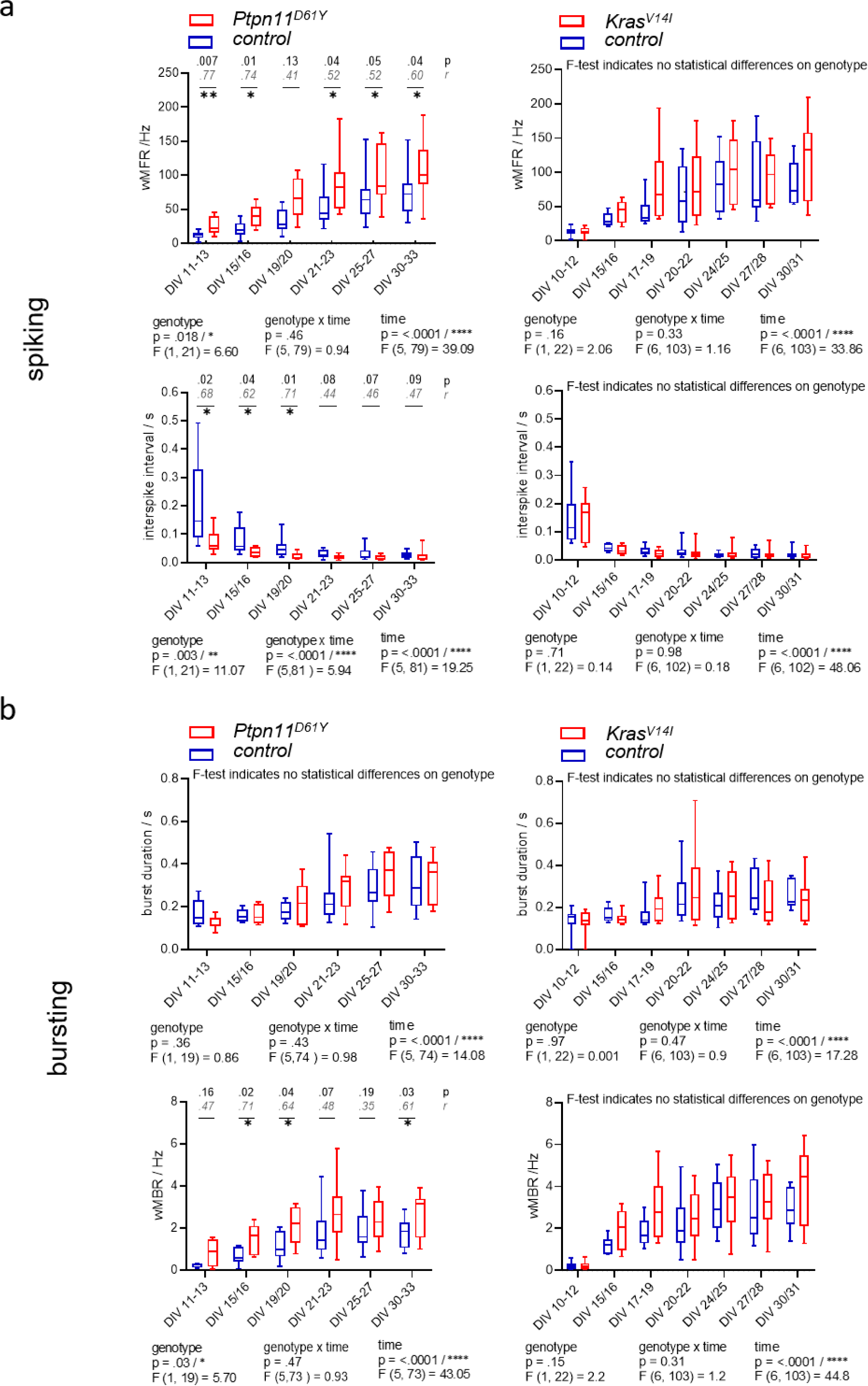
Spontaneous spiking and bursting activity in neuronal networks from the RASopathy models *Ptpn11*^D61Y^ and *Kras*^V14l^. **(a-b)** Parameters describing neuronal activity as a function of culture age as box plots indicating the median and interquartile range with whiskers extending from 5% to 95% confidence interval. **a** wMFR and interspike interval. **b** Burst duration and wMBR. Parameters are statistically evaluated by a mixed-effects model approach. Below the graphs, p and F values for main and intermediate effects are shown. If the F test reveals significance in genotype, p values from the Mann‒Whitney U test (black) and effective size r according to Wendt (gray, cursive) are shown in the graphs. Stars indicate significant differences between genotypes for a time point: < 0.05 (*), < 0.01 (**). A data point represents the mean values across the wells related to one animal/preparation (preparation level).

The projected data set on PC1 was shown to be approximately normally distributed (Supplementary material Figure S 9) with equal variances in all groups (Supplementary material Figure S 10), leading us to perform a mixed-effects model with Sidak’s test to correct for multiple comparisons for increased statistical power. Since concomitant with PCA, normalization of data values centers the data points around zero for each time point, the systematic increase of neuronal activity coming along with network development was filtered out in all groups. Thus, the effect caused by mutation on spiking, bursting and network bursting phenotypes was unveiled, decreasing the probability of error type two. Additionally, for disinhibition-induced stimulation experiments, Sidak’s multiple comparisons test was performed.

To compare the evoked activity upon STIM, we used the nonparametric Mann‒Whitney test due to the nonnormal distribution of the data set for MFR. Additionally, we investigated differences in distribution by the Kolmogorov‒Smirnov test. To analyze the effect of tetanic stimulation on networks, we calculated the relation 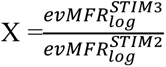 between the logarithmic *evMFR*_*norm*_ after (STIM 3) and prior (STIM 2) tetanic stimulation and checked for alterations between genotypes by Welch’s t test [40]. Therefore, we calculated descriptive measures from the independent, individual experiments first, followed by Welch’s t test according to

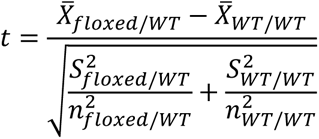

The mean value *X̅* is the average of X across all experiments for the indicated genotype. S^2^ is given by the averaged variance of the indicated genotype derived from the individual experiment, and n is the total number of all wells over all experiments for the indicated genotype. To check the effect of tetanus on networks, we additionally performed a one-sample Wilcoxon test on the pooled data set of X from all experiments.

## Results

### Increase in population-wide neuronal activity in neuronal networks from *Ptpn11*^D61Y^mice

To identify the effect of RASopathy mutations on the functional development of neuronal networks, we performed longitudinal recordings of neuronal network activity in cultured dissociated cortical neurons grown on mwMEA plates. The neurons were derived from the cerebral cortices of newborn mice. As the RASopathy model, we used mice heterozygous for the floxed mutation *Ptpn11*^D61Y^ or *Kras*^V14l^ as well as for the Emx1-cre allele and as the control, their littermates homozygous for the *Ptpn11* or *Kras* wild-type allele and heterozygous for the Emx1-cre allele. The spontaneous network activity of cortical cultures was recorded multiple times in cultures in a time range of DIV 11 until DIV 33 (time series). For one experiment, cultures were prepared from six to eight newborn mice (P0-P1) littermates.

To analyze the parameters describing spontaneous activity in neuronal networks, we averaged the data derived from individual wells across preparation/animal and subsequently grouped the datapoints according to genotype. To assess differences between groups and time points, a mixed-effects model was calculated. Herein, we accepted the violation of the distributional assumption of equal standard deviation within groups due to the robustness of the mixed effects model (Supplementary data Fig. S 8). The global evaluation revealed that all groups reliably exhibited spontaneous activity on DIV 11-13. A gradual increase in population-wide firing activity (assessed as spiking and bursting, Fig. 1a, b) and in functional connectivity/network synchronicity (assessed as network bursting and correlation in spiking, Fig. 2b, c) was evident between DIV 19 and DIV 25, when all parameters stabilized, presumably marking network maturity. Sample sizes are listed in Fig. 2a.

**Fig. 2.**
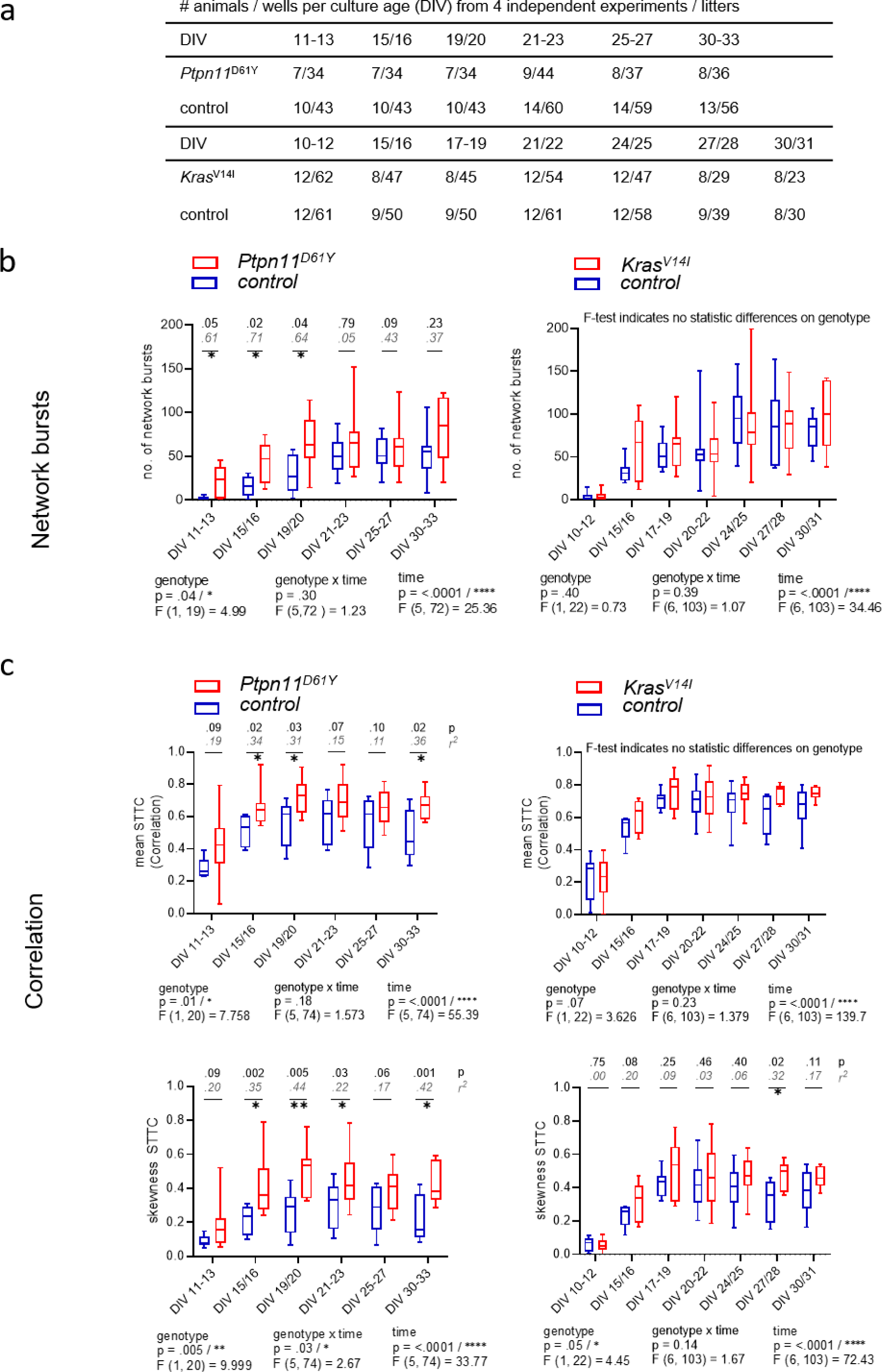
Functional connectivity/synchronicity in networks with RASopathy mutations *Ptpn11*^D61Y^ and *Kras*^V14l^. **a** Number of animals and wells analyzed at each culture age cohort for all parameters of spontaneous activity. **(b-c)** Parameters describing functional connectivity as a function of culture age as box plots indicating the median and interquartile range with whiskers extending from 5% to 95% confidence interval **b** Values of the number of network bursts. **c** Values of mean STTC and skewness STTC. Parameters are statistically evaluated by a mixed-effects model approach. Below the graph, p and F values for main and intermediate effects are shown. If the F test reveals significance in genotype, in **b,** p values from the Mann‒Whitney U test (black) and effective size r according to Wendt (gray, cursive) are shown; in **c,** p values from the unpaired t test together with effect size r^2^ are shown in the graphs. Stars indicate significant differences between genotypes for one time point < 0.05 (*), < 0.01 (**), < 0.001 (***), < 0.0001 (****). A data point represents the mean values across the wells related to one animal/preparation (preparation level).

To assess the effect of RASopathy mutations on spiking and bursting behavior, we followed up on four parameters, including wMFR, interspike interval, burst duration and wMBR (Fig. 1). *Ptpn11*^D61Y^ networks were significantly different in nearly all parameters compared to their control according to the F test in the mixed effects model. The spiking activity was clearly increased as reflected in significantly higher values in wMFR nearly throughout all observation time points (except on DIV 19/20) and in a decreased interspike interval reaching significance at the early points in time (DIV 11-13, DIV 15/16, DIV 19/20) (Fig. 1a, left column). Furthermore, *Ptpn11*^D61Y^ networks showed increased bursting behavior compared to their control (Fig. 1b, left column). The appearance of spontaneous bursts during development is important for the formation of neuronal circuits [20].

*Ptpn11*^D61Y^ networks exhibited significantly higher wMBR on DIV 15/16, DIV 19/20 and DIV 30-33. However, the burst duration remained unchanged. In contrast, we detected no significant differences in any parameter describing spiking and bursting behavior in *Kras*^V14l^. However, visualization of the data indicates a conspicuous trend of higher median values in parameters reflecting neuronal activity (wMFR, wMBR, burst duration) (Fig. 1a, b, right column).

A fundamental feature of neuronal networks is their functional connectivity/network synchronicity, which describes the impact of the activity of other neurons within the same network on the activity of a given neuron [25]. To assess functional connectivity during network development, we calculated the number of network bursts (Fig. 2b) and determined the correlation between all pairs of electrodes during NBs measured as STTC (Fig. 2c). *Ptpn11*^D61Y^ networks globally demonstrated significantly higher functional connectivity than their control groups, as revealed by a mixed effects model in all observed parameters (Fig. 2b, c, left side). In *Ptpn11*^D61Y^ networks, the number of NBs was increased at early time points (DIV 11-13, DIV 15/16 and DIV 19/20), but this effect was abolished at later time points. Moreover, compared to controls, *Ptpn11*^D61Y^ networks indicated significant differences in the mean and skewness of STTC throughout the whole observation time, as tested by multiple comparison tests (Figure 2C, left side). In contrast, compared to the control group, *Kras*^V14l^ showed no alterations in the number of NBs or in the mean STTC (Fig. 2b, c, right side). However, a significant global difference was calculated for the skewness of STTC according to the F test on the main effect genotype on DIV 27/28. In general, we identified elevated predicted median values in the *Kras*^V14l^ networks in most parameters; however, the differences were not significant, likely due to large variability, which is typical of MEA-derived data.

### Principal component analysis of MEA parameters reveals differences in genotype clusters and identifies differences in spiking, bursting and network bursting in both RASopathy models

Analysis of most individual parameters from the longitudinal recordings in *Ptpn11*^D61Y^ neurons revealed higher effect sizes in younger cultures that were reduced at later time points. Similar effects were observed in *Kras*^V14l^ neurons, although not significantly. To increase the interpretability of data sets and the power of statistical testing and to allow simpler visualization, we introduced PCA, which reduces data dimensionality. We performed PCA including 19 parameters obtained from a custom-written MATLAB program (Supplementary material Table S 2). We conducted the PCA at the well level. In contrast to the preparation level, we yielded a higher number of data points, resulting in a lower probability of error type two but decreased power.

We analyzed all age cohorts from the longitudinal recordings and visualized the transformed data points in a PC1 and PC2 coordination system (Fig. 3a, Fig. 4a). Sample numbers are shown in Fig. 3d and Fig. 4d. In both RASopathy disease models, the respective clusters and their controls appear to converge with culture age, also indicated by the visualized distance between the cluster centroids in the graphs. To quantify the separation between genotype clusters, we calculated the Mahalanobis distance that accounts for the correlation of the variables [24]. Statistical significance was evaluated by Hotelling’s two-sample T^2^ related to an F value as introduced earlier [13]. For both RASopathy model networks, the Mahalanobis distance to the control group turned out to be strongly significant at the early stages but dramatically decreased with culture age (Fig. 3b, Fig. 4b). At the end of the observation period, *Ptpn11*^D61Y^ networks still differed significantly from their control group according to the F-statistic on DIV 30-33, while for Kras^V14l,^ the clusters converged more closely and did not differ significantly on DIV 27/28. The reliability of the results is underpinned by the cumulative fraction of variance of PC1 and PC2 explaining approximately 70% of the variation in both data sets (Fig. 3c, Fig. 4c). This approach confirmed a developmental phenotype in spontaneous network activity in both RASopathy disease models.

**Fig. 3.**
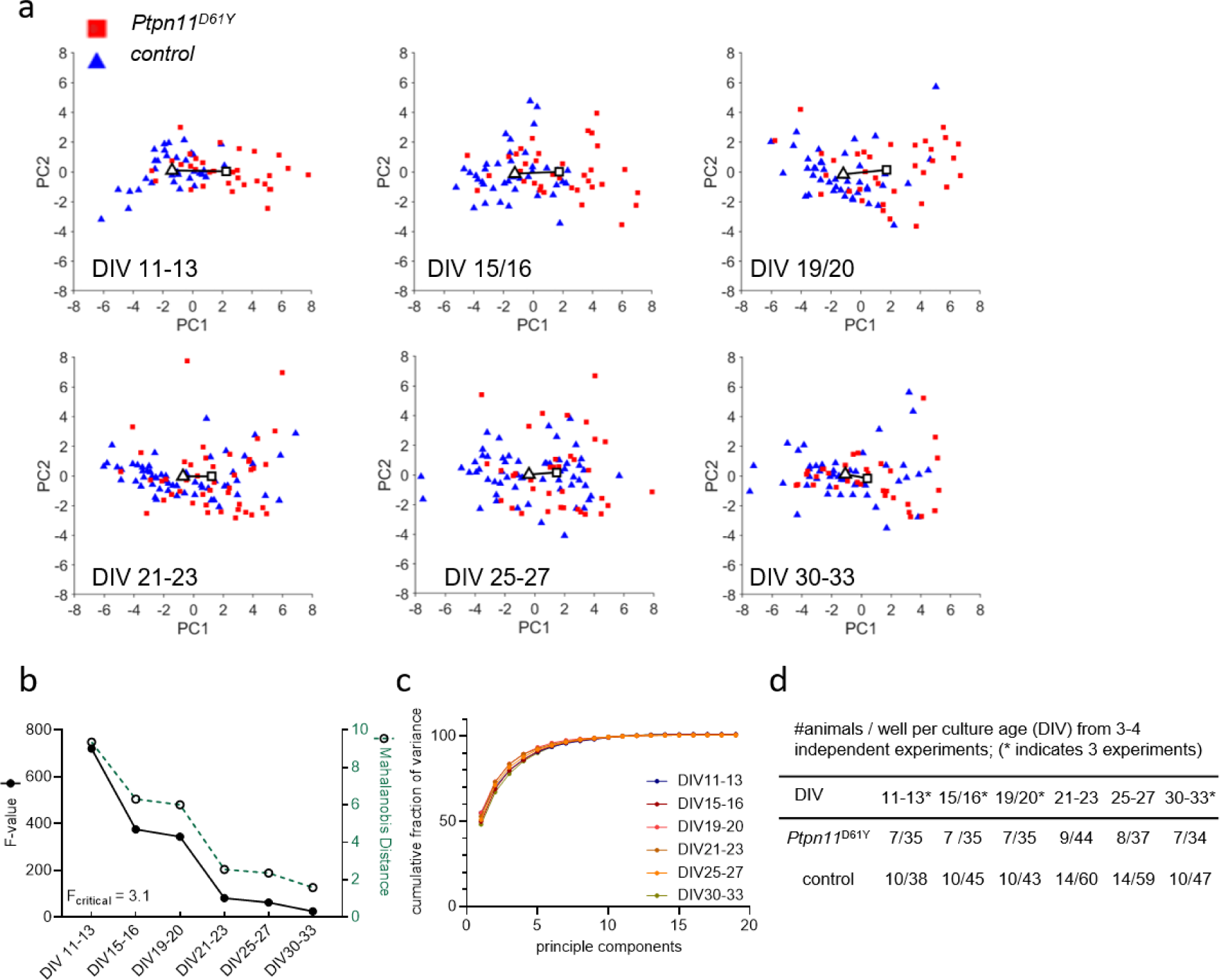
PCA of the *Ptpn11*^D61Y^ 19-dimensional parameter vector. **a** Principal component PC1 versus PC2 scores plotted as a function of culture age (DIV). A solid line is drawn between the centroids of the control cluster (triangle) and mutant cluster (rectangle). Each color-coded data point represents a recording from one well. **b** Cluster separation in dependence of culture age was measured by Mahalanobis distance (green dotted line), and the F value shows statistical significance (black solid line) (F_critical_ = 3.1). **c** Fraction of variance explained by each PC summed over all PCs. The cumulative fraction of variance is indicated for each evaluated culture age. **d** Sample size is given per culture age as the number of animals and wells.

**Fig. 4.**
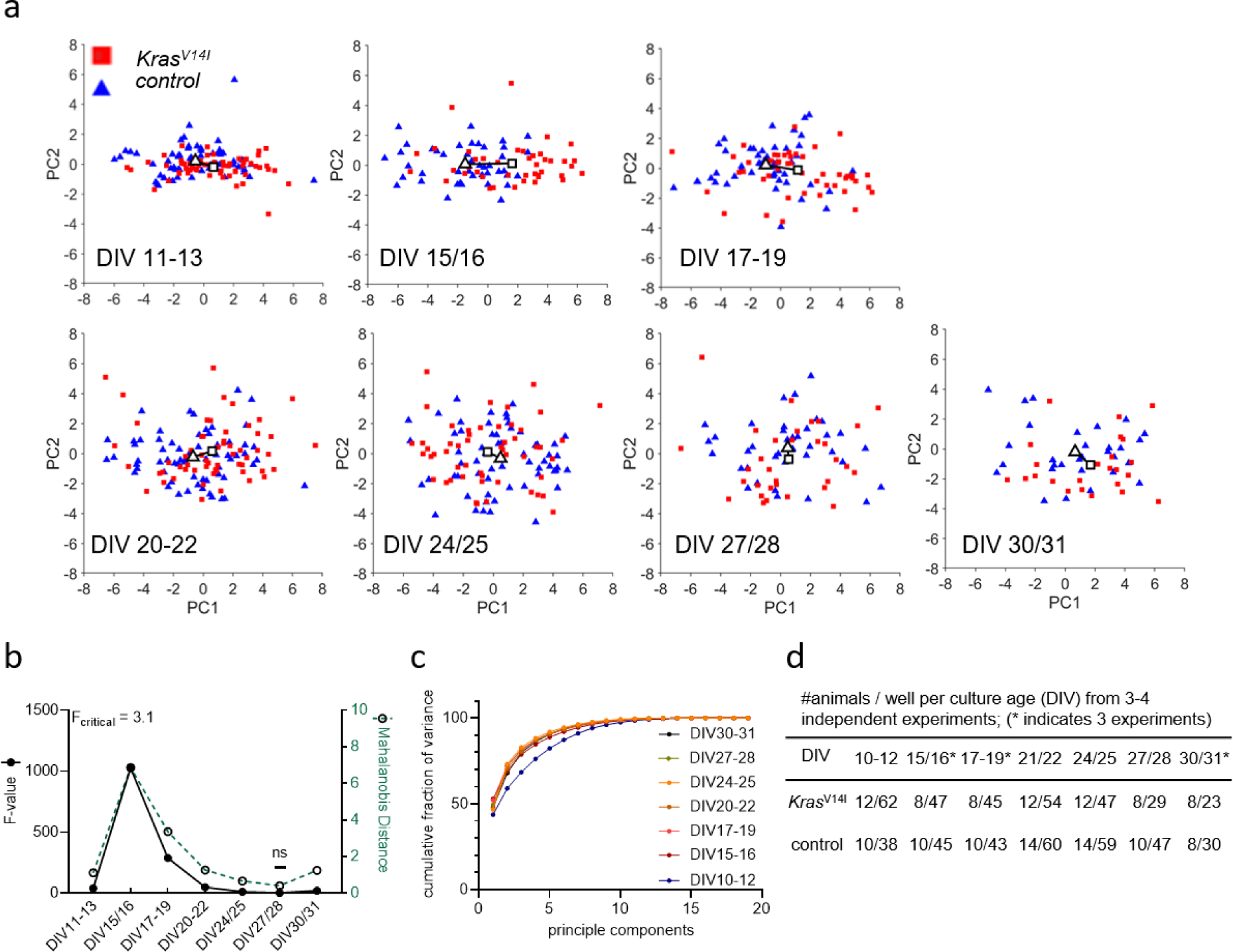
PCA of the *Kras*^V14l^ 19-dimensional parameter vector. **a** Principal component PC1 versus PC2 scores plotted as a function of culture age (DIV). A solid line is drawn between the centroids of the control cluster (triangle) and mutant cluster (rectangle). Each color-coded data point represents a recording from one well. **b** Cluster separation in dependence of culture age was measured by Mahalanobis distance (green dotted line) between the clusters of genotypes, and the F value shows statistical significance (black solid line) (F_critical_ = 3.1). **c** Fraction of variance explained by each PC summed over all PCs. The cumulative fraction of variance is indicated for each evaluated culture age. **d** Sample size is given per culture age as the number of animals and wells.

To further identify at which level spontaneous network activity is particularly affected in RASopathy models, we performed a further PCA that specifically combines parameters describing the features of spiking, bursting and network bursting behavior (Supplement Material Fig. S5b). Therefore, we formed parameter vectors from the extracted parameters from the spontaneous activity analysis. The values were logarithmized for nonnormally distributed data. This resulted in a 4-dimensional vector for spiking (log_10_(wMFR), log_10_(wtMFR), log_10_(interspike interval), and number of contributing electrodes), a 3-dimensional vector for bursting (log_10_(wMBR), log_10_(burst duration), and log_10_(number of spikes per burst)) and a 4-dimensional parameter vector for network bursting (number of network bursts, mean NB duration, number of spikes per NB, and number of contributing channels). The PCA resulted in a component projection on PC1 for each age cohort (Fig. 5). We visualized the projected parameter vector on PC1 for each neuronal age cohort (Fig. 5a-c). We were able to identify a changed phenotype in spiking, bursting and network bursting for both RASopathy models compared to wild type.

**Fig. 5.**
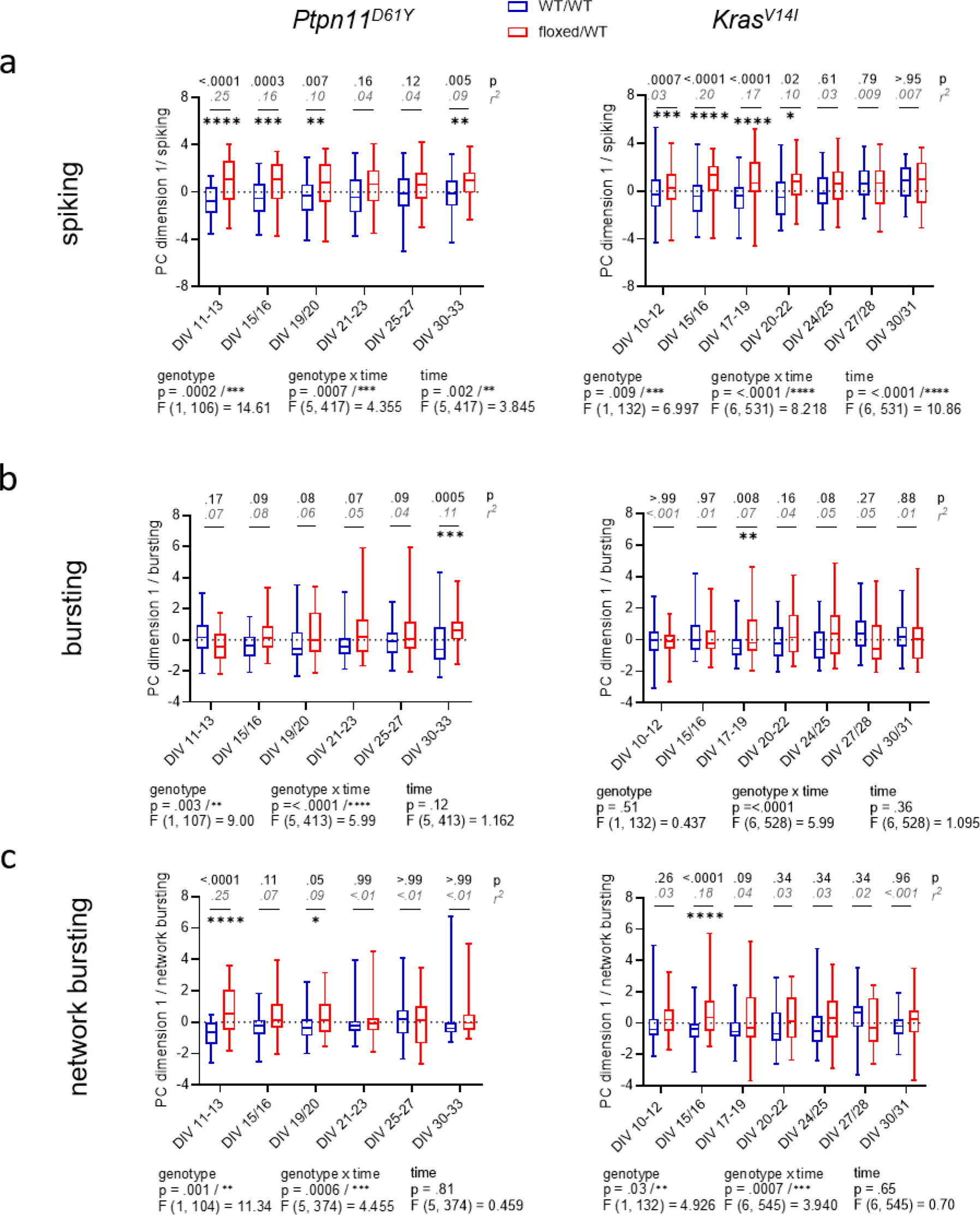
PCA of parameter vectors related to spiking, bursting and network bursting from *Ptpn11*^D61Y^ and *Kras*^V14l^ **(a-c)** PC1 projection as a function of culture age as box plots indicating the median and interquartile range with whiskers extending from 5% to 95% confidence interval. **a** Quantification of spiking behavior calculated as a projection of a 4-dimensional parameter vector (number of spiking electrodes, wMFR, wtMFR, interspike interval). **b** Quantification of bursting behavior calculated as the projection of a 3-dimensional parameter vector (wMBR, bursting duration, no. of spikes per burst). **c** Quantification of network bursting calculated as a projection of a 4-dimensional parameter vector (number of network bursts (NB), mean NB duration, number of spikes per NB, mean number of contributing channels per NB). Statistical significance was assessed by a mixed-effects model approach. p values (in black) for each comparison were obtained by Sidak’s multiple comparisons test, and effect size (r^2^, in gray, cursive) was derived from the unpaired t test. Stars indicate significant differences in median values between genotypes for time points:.05 (*),.01 (**),.001(***), <.0001(****).

On the spiking level, the mixed effects model revealed a strong significance between the group mean in the main effect genotype (Fig. 5a). Therefore, multiple comparisons testing showed that the strong differences were primarily at early time points (*Ptpn11*^D61Y^: DIV 11-13, DIV 15/16, DIV 19/20; Kras^V14l^: DIV 15/16, DIV 17-19, DIV 20-22). Similarly, for network bursting, the mixed effects model indicated a significant difference in genotype with strong differences on DIV 11-13 and on DIV 15/16 in *Ptpn11*^D61Y^ and *Kras*^V14l^, respectively (Fig. 5c). In contrast, as assessed by multiple comparisons testing, for bursting, only *Ptpn11*^D61Y^ showed significant differences at later time points (DIV 21-23, DIV 30-33), when networks are expected to be mature. Interestingly, the mixed effects model identified a strong interaction effect over time and between genotypes in all parameter vectors describing the network activity for both RASopathy models (genotype x time, indicated in all graphs in Fig 5).

Overall, we particularly observed an alteration in spiking and network bursting for both RASopathy models during network development, but there were also effects on bursting caused by RASopathy mutation. Data also indicate that differences are more pronounced during early development and shades out during network maturation.

### Pharmacological disinhibition has a specific effect on network bursting in *Ptpn11*^D61Y^

Our experiments revealed a change in neuronal activity and functional connectivity during network development in both RASopathy models. Interestingly, the effect of mutations on network characteristics is attenuated over time, pointing to possible adaptation.

network activity. Therefore, we applied bic, a potent antagonist of the GABA_A_ receptor, to cultured cortical neurons (DIV 33) from *Ptpn11*^D61Y^ ^mice^ and their wild-type littermates.

As expected, the bic treatment resulted in a reliable increase in spiking activity reflected in increased wMFR (Figure 6b, c), which was statistically confirmed by one sample Wilcoxon test for each well (Figure 6c). While spiking and bursting behavior did not differ between genotypes upon bic treatment (Fig. 6c, Supplementary Material Fig. S 11), we detected elevated network bursting in disinhibited *Ptpn11*^D61Y^ networks compared to the control. The mixed effects model revealed a global interaction effect of time and genotype (genotype x time p <.0001), leading us to perform Sidak’s multicomparison tests resulting in significant differences from minute 50 post bic (Fig. 6c, d). For an overall conclusion of the main effect genotype, the mixed effects model returns a p value of.06 that is near significance. This result revealed changes in the inhibitory system in *Ptpn11*^D61Y^ networks, which likely developed to counteract increased spontaneous network activity.

**Fig. 6.**
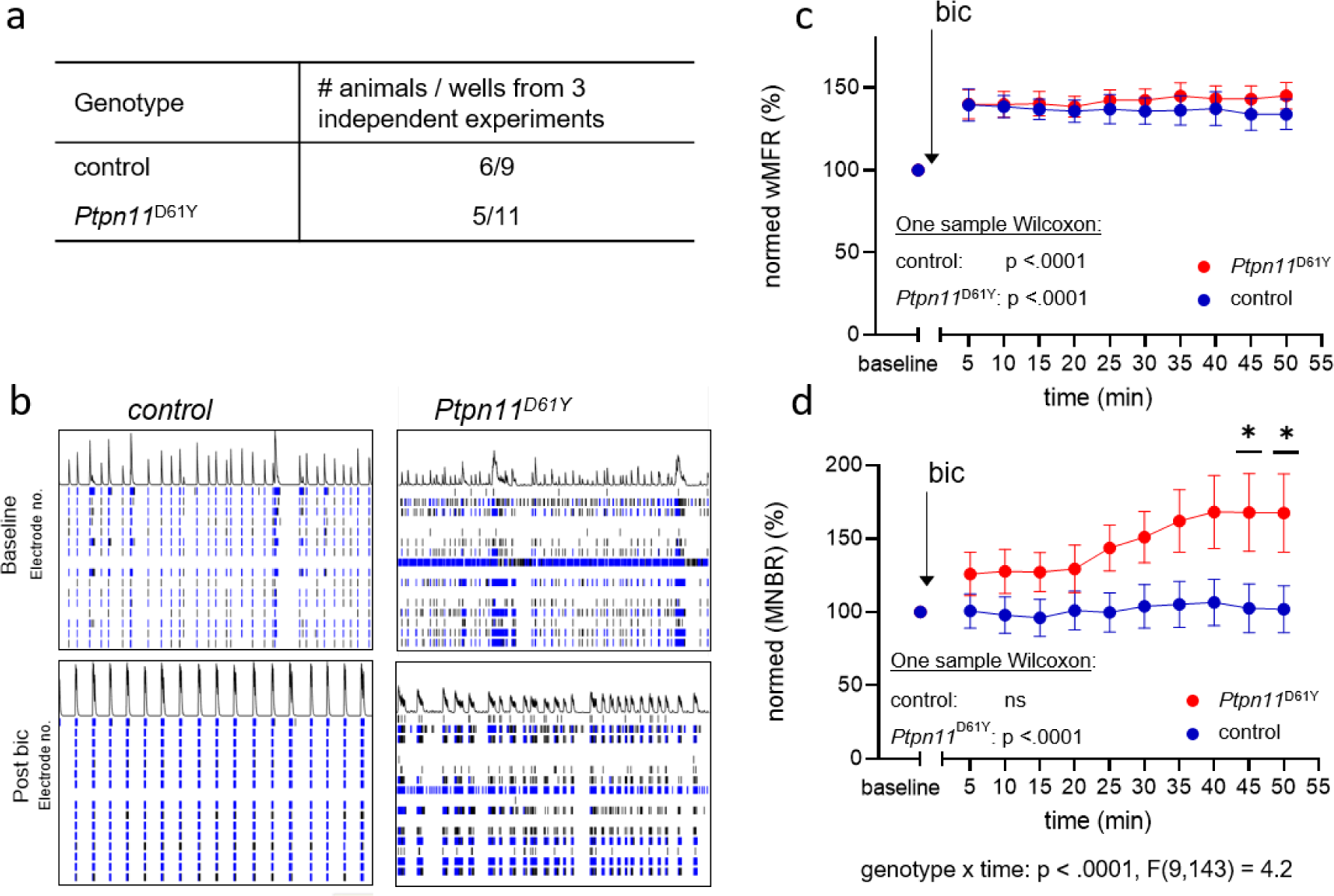
Effect of disinhibition on neuronal network activity in *Ptpn11*^D61Y^ on DIV 33. **a** Number of animals (from 3 independent experiments) and wells used for this analysis. **b** Representative raster plots showing spatiotemporal spiking activity of control and *Ptpn11*^D61Y^ networks prior to and 30 min after bicuculline application (bic). (**c-d)** Line graphs of wMFR **(c)** and MNBR **(d)** as a function of time prior to and after application of bic showing an elevated spiking upon treatment in both genotypes. Values are represented in percent relative to a baseline generated from values recorded over a time interval of 20 min prior to bic treatment, averaged and set to 100%. A significant increase against baseline was statistically tested by one-sample Wilcoxon. In **(d)**, the interaction effect between time and genotype was calculated by a mixed effects model approach. Sidak’s multiple comparison tests were used to determine significance between genotypes. Stars indicate significant differences < 0.05 (*). To assess the possible contribution of a shift in excitatory/inhibitory (E/I) balance, as previously suggested in [28], we decided to test the effect of pharmacological interference with inhibitory tone on

### Fewer GABAergic synapses but higher expression of vesicular transporters for GABA and glutamate were detected in the networks with the *Ptpn11*^D61Y^ mutation

To examine the altered inhibitory tone on a synaptic level, we visualized and quantified glutamatergic and GABAergic boutons in *Ptpn11*^D61Y^ networks and their control. Therefore, neuronal cultures were immunolabeled with specific antibodies against vesicular GABA transporter (VGAT) or vesicular glutamate transporter (VGLUT1), together with antibodies against synaptophysin (Syp), to visualize all synapses (Fig. 7a, b). We used our in-house developed MATLAB-based program SynEval to obtain Syp-positive puncta (ROIs) and to sort them as VGAT- or VGLUT-positive in an automatized batch mode, which reduces human bias and time effort. The fraction of glutamatergic boutons was significantly higher in the *Ptpn11*^D61Y^ group than in the control group (Fig. 7d), and accordingly, the fraction of GABAergic boutons decreased (Fig. 7c). Interestingly, we noticed an increased synaptic IFI for VGAT or VGLUT, indicating increased synaptic abundance of these neurotransmitter transporters (Fig. 7c, d). These findings further point toward changes in the excitation-inhibition balance in *Ptpn11*^D61Y^ networks.

**Fig. 7.**
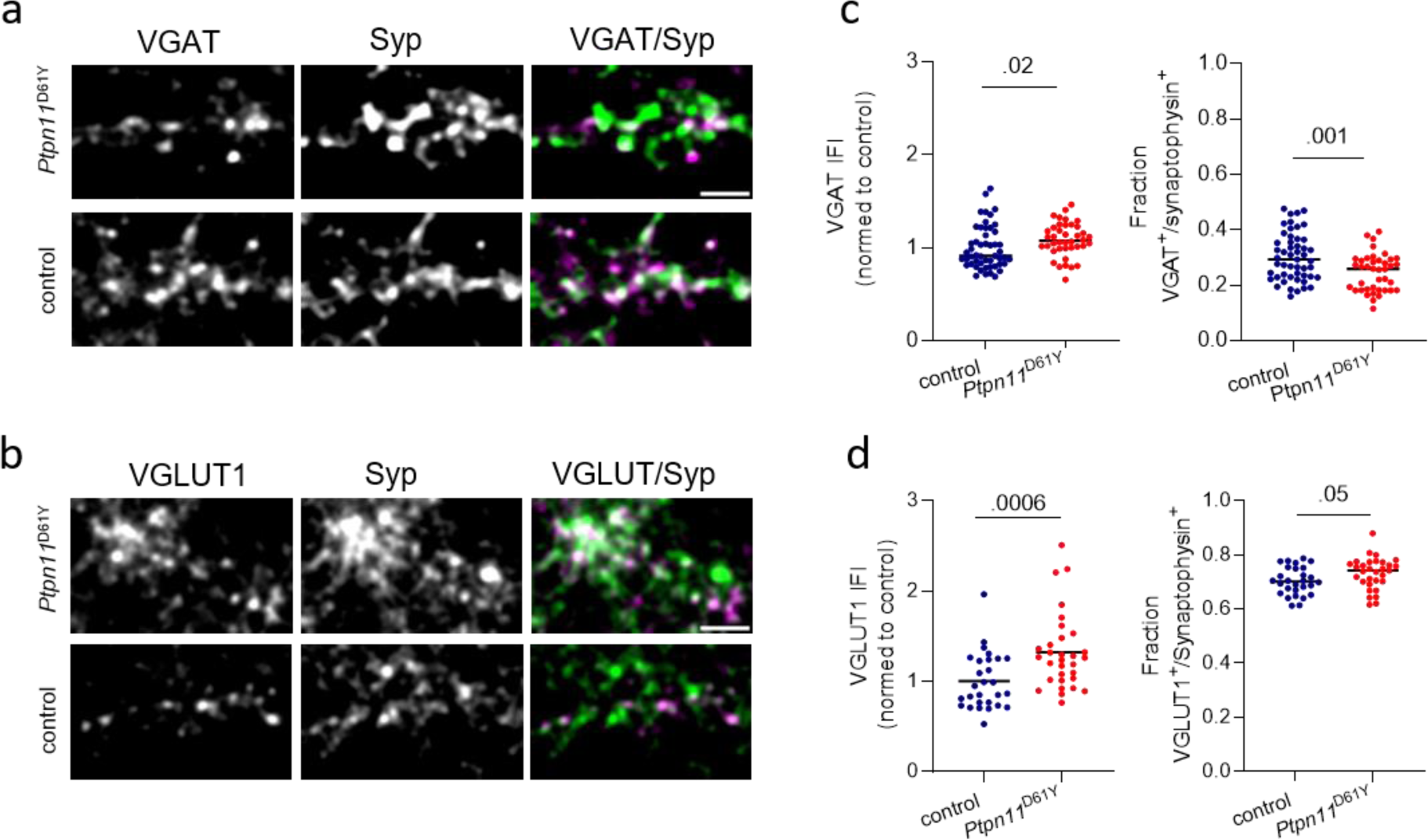
Immunolabeling of excitatory and inhibitory synapses in neuronal networks from *Ptpn11*^D61Y^ mice. **a** Representative images of neurons (DIV 21) labeled using antibodies against VGAT (green) and Syp (red). **b** Representative images of neurons labeled using antibodies against VGLUT1 (green) and synaptophysin (Syp, red). **c** Quantification of VGAT IFI in Syp-positive synaptic puncta and of the fraction of VGAT-positive synapses. **d** Quantification of VGLUT1 IFI in Syp-positive synaptic puncta and of the fraction of VGLUT1-positive synapses. Statistical significance was determined by the Mann‒Whitney test. Data points correspond to individual analyzed visual fields derived from two independent experiments. The scale bar is 5 µM.

### Altered evoked activity in *Ptpn11*^D61Y^ cells probed by electrical stimulation

The longitudinal analysis indicated a normalization of spontaneous network activity during maturation in the neuronal networks expressing RASopathy mutations. To assess whether there are functional defects in the processing of information in these apparently “normal” networks, we analyzed network activity evoked by the delivery of electrical stimulation. We recorded network activity evoked by single-pulse stimulation (test stimulus, STIM) delivered from different electrodes in *Ptpn11*^D61Y^ networks grown for 33 days (DIV 33) (Supplement Material Fig. S 3a, d). Of note, we solely analyzed robust and stable networks, which were not affected by the application of STIM itself, i.e., the activity evoked by STIM1 was the same as the activity evoked by STIM2 (cf. Material and Methods, Data cleaning and selection of valid wells). The data points plotted in Fig. 8a display the evMFR_norm_ representing the evoked MFR induced by STIM at the electrode level and normalized to recorded spontaneous activity on the same electrode, averaged across an array (for detailed information on calculation, see Supplementary Material Fig. S 6). Interestingly, evMFR_norm_ was found to be significantly decreased in *Ptpn11*^D61Y^ networks compared to wild type (Fig. 8a, upper plot). The graph shows the data for STIM2, and similar results were obtained for STIM1 (Supplementary Material Fig. S12). The Kolmogorov‒Smirnov test confirmed a different distribution of data between both genotypes. The dampened electrical excitability of *Ptpn11*^D61Y^ networks is in line with our finding of affected inhibitory tone and the decrease in synaptic activity in *Ptpn11* ^D61Y^ shown in [3].

**Fig. 8.**
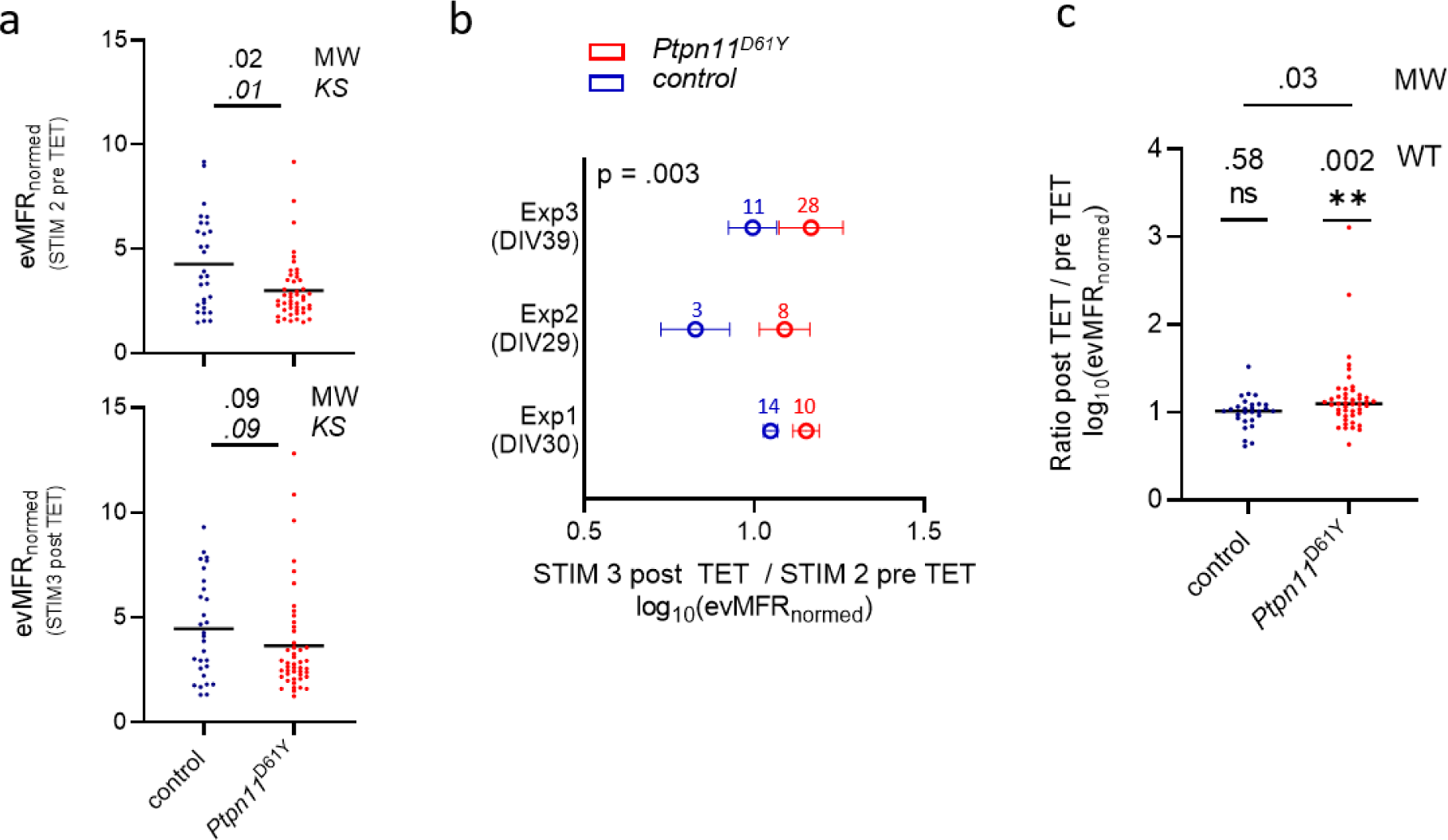
Assessment of evoked network activity in *Ptpn11*^D61Y^ neurons. **a** Evoked mean firing rate (evMFR_normed_) before and after TET stimulation. Black lines indicate median values, and data points correspond to one well. Significance was tested by the Mann‒Whitney (MW) test to check differences in mean values and the Kolmogorov‒Smirnov (KS) test to check differences in data distributions. **b** Relative change (ratio) of evMFR_normed_ recorded before and after application of TET are shown from three independent experiments (Exp1-Exp3). Whiskers indicate SEM. The p value was calculated by Welch’s t test; the number of included wells for the corresponding genotype is color-coded for each experiment. **c** Analysis of data pooled from all 3 experiments. Black lines indicate median values, and one data point refers to one well. The one-sample Wilcoxon test (WT) was tested against a hypothetical value of 1 (indicating no change in evMFR_normed_ caused by TET), and the Mann‒Whitney test (MW) was used to calculate significant differences between genotypes.

### Effect of high-frequency electrical stimulation on evoked activity in *Ptpn11*^D61Y^ mice

We observed that evoked neuronal activity upon STIM is potentially affected by RASopathy-associated mutations and that excitability is dampened. Next, to understand whether neuronal plasticity might also be affected in *Ptpn11*^D61Y^ networks, we tested the effect of tetanic stimulus (TET) on evoked synaptic activity. The stimulation protocol described in Supplementary Material Fig. S 3c was previously suggested to be suitable for analysis of neuronal plasticity since it affected evoked activity in networks [9]. The TET stimulus applied on *Ptpn11* ^D61Y^ networks induced different effects compared to controls in all three independent experiments individually plotted in Fig. 8b. Calculating the relative change (ratio) between post and pre TET logarithmic (log_10_)evMFR_norm_ showed that the mean value in *Ptpn11*^D61Y^ was higher than that in the control group for each experiment (Exp1-Exp3), with an overall significant difference confirmed by Welch’s t test (Mann‒Whitney test for individual experiments: Exp1: p = 0.03, Exp2: ns, Exp3: ns). More precisely, the findings reveal that TET resulted in a significant increase in evMFR_norm_ in *Ptpn11*^D61Y^ networks while not inducing an effect in controls (Fig. 8c). That was shown by testing the pooled data set against the hypothetical value 1 (no change between pre and post TET logarithmic *evMFR*_*norm*_) by a one-sample Wilcoxon test. Interestingly, the difference in evMFR_norm_, demonstrated between *Ptpn11*^D61Y^ and control pre TET, was diminished post TET (Fig. 8a, bottom plot). According to the Kolmogorov‒Smirnov test, the difference in distribution also failed significance after TET. Indeed, in contrast to the control, *Ptpn11*^D61Y^ demonstrated a shift between STIM2 and STIM3 curves in their relative frequency plots, particularly for higher *evMFR*_*norm*_, pointing to an altered distribution in the data set (Supplementary Material Fig. S12b, c).

## Discussion

RASopathies are a group of rare genetic disorders with various symptoms, including heart defects, skeletal anomalies, increased risk for tumors and variable degrees of neurocognitive deficits. Clinically, the severity of neurocognitive impairments in RASopathies often varies over the lifetime and often becomes milder with the age of the patients. Neurodevelopmental phenotypes were also described in RASopathy mouse models, where the involvement of maladaptive changes in cortical networks was suggested [28].

To investigate whether similar maladaptation occurs in neuronal networks on a chip, we cultured cortical neurons carrying two distinct RASopathy mutations on mwMEAs and recorded their population-wide spontaneous network activity to monitor network establishment and maturation during a time period of 5 weeks. To facilitate the evaluation of large data sets acquired longitudinally with a high sampling rate (12.5 kHz), we developed and implemented an analysis pipeline programmed in MATLAB. From the list of spikes per electrode, provided by acquisition software, it computes a custom set of parameters describing various features of neuronal spontaneous network activity, including spiking, bursting, network bursting and correlation in spiking. This custom-written software package permits fast and unbiased analysis of large MEA data sets while being flexibly applicable for a wide set of experimental designs in general. The provision of a GUI renders the package user-friendly. Furthermore, we developed an add-on package of MATLAB-based functions that implements PCA to reduce the complexity of analysis outcomes by projecting the resulting parameters into a two-dimensional system. The package allows the calculation of Mahalanobis distance and statistical testing to mathematically compare the network behavior between two groups derived in our case from mutant and wild-type animals and determine if these constitute significantly distinguishable populations. In the next step, it also computes whether the difference between network clusters can contribute to one specific network function feature (spiking, bursting and network synchronicity) or whether changes in multiple features contribute to the clusters’ differences. This approach turned out to be suitable to uncover effects in the MEA data that are inherently characterized by high complexity and variability.

On the basis of MEA-recorded data and supported by the MATLAB analysis pipeline, we identified a convergent developmental phenotype in both RASopathy models. Multiple parameters describing neuronal activity and functional connectivity were significantly different in neurons with the *Ptpn11*^D61Y^ mutation. The differences were more pronounced in younger networks and faded out as networks matured. In contrast, the neurons with the *Kras*^V14l^ mutation showed only a tendency toward similar changes in spontaneous network activity parameters as seen in *Ptpn11*^D61Y^ neurons. Being aware of the high inherent variance of MEA data, we performed PCA to obtain an overall description of network parameters for both models. This analysis revealed a significant separation between clusters representing neurons with the *Ptpn11*^D61Y^ or *Kras*^V14l^ mutation and their respective controls. Moreover, for both mutations, the cluster separation from control networks decreased throughout development, and consecutive analysis identified a major contribution of changes in spiking and network bursting to the PCA cluster separation. Thus, based on the analysis of spontaneous network activity, we have clearly demonstrated similar developmental phenotypes for both mutations, confirming that the effect of RASopathy mutations on neuronal network function converges. It is interesting to note that mutations’ effect on network function correlates with the strength of their activation assessed biochemically in previous studies. While *Kras*^V14l^ exhibits attenuated biochemical activation compared to common cancer-associated KRAS alleles [56], strong activation was shown for *Ptpn11*^D61Y^, which is even linked to juvenile myelomonocytic leukemia (JMML) in patients [38].

The normalization of network activity parameters in mature networks suggests that compensatory mechanisms develop to counteract the increased network activity observed in juvenile networks. A shift in the E/I balance might represent one such mechanism. Indeed, the balance between E/I within neuronal circuits is crucial for correct brain development and function, and its disturbances contribute to numerous neurodevelopmental disorders, such as autism spectrum disorders and epilepsy, which are common comorbidities in more severe RASopathies, including CFC or CS [1, 30]. Moreover, the recent characterization of a RASopathy animal model expressing the strongly activating allele *Kras*^G12V^ indicated that increased inhibitory drive contributes to the electrophysiological and behavioral abnormalities in this model [28]. In line with these assumptions, we demonstrated increased network bursting in mature *Ptpn11*^D61Y^ networks upon disinhibition mediated by pharmacological blockade of GABA receptors. The immunocytochemical analysis revealed an increased proportion of excitatory and decreased proportion of inhibitory synapses in *Ptpn11*^D61Y^ networks as well as an increased expression of vesicular transporters for both glutamate and GABA. In line with a modified inhibitory tone, we found that the application of electrical test stimuli at a low frequency elicited a reduced evoked response in the *Ptpn11*^D61Y^ networks, pointing toward maladaptive dampening of network activity, which leads to an attenuated response to stimulation. An application of TET stimulation that had no effect on evoked responses in the control had a significant impact on responses in neurons with the *Ptpn11*^D61Y^ mutation. Interestingly, TET stimulation statistically restored the affected evoked activity in the RASopathy model. However, the change in the distribution of data values between pre- and posttetanus stimulations indicates that there is much higher variability in the responses of individual tested networks expressing *Ptpn11*^D61Y^ mutations compared to controls. At this timepoint, it is not clear which concrete parameters are involved in RASopathy-induced network adaptation. A better understanding of the circumstances under which plasticity can be induced can potentially be used in therapeutic approaches to break the maladaptive circuits and use them for targeted treatment of RASopathies and in their comorbidities.

In summary, our study reveals that different RASopathy mutations have convergent effects on neuronal network activity. We observed more severe differences in network characteristics in juvenile networks that were compensated in the late stages of network maturation, which is in good agreement with clinical data. Despite the apparent normalization of spontaneous network activity, we uncovered significant differences in E/I balance and in evoked responses in mature networks. Finally, the cell on-a-chip approach developed here is well scalable, accessible for treatments and manipulations and adaptable for stem cell technology-derived humanized disease modeling. Therefore, we assume that the analysis workflow developed here could also be implemented by drug screening and testing platforms for unrelated brain diseases.

## Supporting information

Supplementary Material Complete

## Abbreviations

AxIS: Axion Integrated Studio
bic: bicuculline
CFC: cardio-facio-cutaneous syndrome
CS: costello syndrome
DIV: days in vitro
e_act_: currently active number of active electrodes
e_max_: maximum number of active electrodes
E/I: excitation and inhibition
FCS: fetal calf serum
GUI: graphical user interphase
ICC: Immunocytochemistry
IFI: immunofluorescence intensity
JMML: juvenile myelomonocytic leukemia
MAPK: mitogen-activated protein kinase
MBR: mean bursting rate
MEA: multielectrode array
MFR: mean firing rate
mwMEA: multiwell MEA
NB: network burst
NBA: Neurobasal^TM^ -A media
NS: Noonan syndrome
PC: principle component
PCA: Principal component analysis
PCR: polymerase chain reaction
ROI: region of interest
RT: room temperature
SA: spontaneous activity
SD: standard deviation
STIM: test stimulation
STTC: spike time tiling coefficient
TET: tetanus stimulation
VGAT: vesicular GABA transporter
VGLUT1: vesicular glutamate transporter
wMBR: weighted mean bursting rate
wtMFR: mean firing rate weighted to total number of electrodes

## Declarations

### Ethics approval and consent to participate

Not applicable

### Consent for publication

Not applicable

### Availability of code, data and material

All custom written source code used for analyzing experiments and plotting is available on a GitHub repository. Code for analyzing spontaneous network activity by functional analysis of neuronal activity and connectivity can be found at Analysis of spontaneous network activity, functions to perform PCA, Mahalanobis distance calculation and corresponding statistics at Principle Component Analysis and for analyzing electrically evoked activity at Analysis of electrically evoked activity.

Data were obtained according to the material section. The data sets during and/or analyzed during the current study are available from the corresponding author upon reasonable request.

### Authors contribution

EW and AF: conception and design of the study, interpretation of data, writing of the original manuscript. EW, DG, AP, VH: data acquisition. EW: development of new methodology and software, data analysis. EW, AP: data visualization. MZ: provided transgenic mice. EW, MZ, AF funding acquisition. All authors have read and approved the final manuscript.

## Acknowledgment

We are deeply grateful to all those who contributed to the success of this research project. First, we would like to thank our cooperation partner from the stem cell biology department at the University Hospital Erlangen, particularly Prof. Dr. Beate Winner, Dr. Tania Rizo and Michaela Farrell for great technical support. Furthermore, we thank the group of Prof. Dr. Thomas Winkler at the Department of Biology at Friedrich-Alexander University of Erlangen for their support.

## Disclaimer

Funded by the European Union. Views and opinions expressed are, however, those of the author(s) only and do not necessarily reflect those of the European Union or the Health and Digital Executive Agency. Neither the European Union nor the granting authority can be held responsible for them.

