## Supplementary Material Complete for "Developmental effect of RASopathy mutations on neuronal network activity on a chip"

### Supplementary Tables

**Table S 1** Chemical reagents and antibodies

| **Reagent** | **Source** | **Identifier** |
|---|---|---|
| Neurobasal™ -A, minus phenol red | Gibco™, Thermo Fisher Scientific | Cat#12349-015 |
| GlutaMAX™ Supplement | Gibco™, Thermo Fisher Scientific | Cat#35050-038 |
| B-27™ Supplement (50X), serum free | Gibco™, Thermo Fisher Scientific | Cat#17504-044 |
| Sodium Pyruvate (100 mM) | Gibco™, Thermo Fisher Scientific | Cat#11360-070;<br>CAS 113-24-6 |
| Antibiotic-Antimycotic (100X) | Gibco™, Thermo Fisher Scientific | Cat#15240-062 |
| HBSS without Mg/Ca (HBSS--) | Gibco™, Thermo Fisher Scientific | Cat#14175-129 |
| Papain Dissociation System (PDS) Kit, EBSS Vial | Worthington Biochemical, Lakewood, NJ, USA | Cat# LK003188 |
| Desoxyribonuclease I | Worthington Biochemical, Lakewood, NJ, USA | Cat# LS0002139 |
| Dispase II | Supply Solutions Roche | Cat# 4942078001<br>CAS 42613-33-2 |
| Poly-L-lysine hydrobromide | Sigma Aldrich | Cat# P1524<br>CAS 25988-63-0 |
| (+)-Bicuculline | Tocris Bioscience, Bristol, UK | Cat# 0130/50<br>CAS 485-49-4 |
| FCS (fetal bovine serum) | Sigma-Aldrich | Cat#F9665 |
| Fluoroshield without DAPI | Sigma-Aldrich | Cat#F6182 |

| **Antibodies** | **Source** | **Dilution** | **Identifier** |
|---|---|---|---|
| rabbit antibodies against VGAT cytoplasmic domain | Synaptic Systems | 1:1000 | RRID#AB_887869, Cat#131003 |
| rabbit antibodies against VGLUT1 | Synaptic Systems | 1:1000 | RRID#AB_887875, Cat#135303 |
| Fluorescent secondary antibodies anti-rabbit Alexa Fluor 488 | Jackson Immunoresearch | 1:1000 | RRID#AB_2313584, Cat#711-545-152 |

| | | | |
|---|---|---|---|
| Fluorescent secondary antibodies anti guinea-pig Cy5 | Jackson Immunoresearch | 1:1000 | RRID#AB_2340462, Cat#706-175-148 |

**Table S 2** Features that were used in the principle component analysis and a brief description about their calculation

| Feature | Description |
|---|---|
| mean firing rate (MFR) | Number of spikes detected in individual arrays during recording time divided by recording time |
| weighted mean firing rate to total (t) electrode number (wtMFR) | MFR weighted by the relation of currently active electrodes to number of electrodes (herein 16) |
| weighted mean firing rate (wMFR) | MFR weighted by the relation of currently active electrodes to number of active electrodes to reference point (e.g. time of maturity in time series) |
| interspike interval | Mean time between spikes in individual arrays during recording time |
| CV interspike interval | Coefficient of variance of ISI in individual arrays during recording time |
| Weighted mean bursting rate (wMBR) | The mean number of bursts in individual arrays per time weighted by the relation of currently active electrodes to number of active electrodes to reference point (e.g. time of maturity in time series), (Hz) |
| Burst duration | Mean duration of bursts calculated as the time from the first spikes to the last spike in individual arrays. |
| CV burst duration | Coefficient of variance of burst duration in individual arrays |
| Mean number of spikes per burst | The average number of within-burst spikes in individual arrays during recording time |
| CV number of spikes per burst | Coefficient of variance of number of within-burst spikes in individual arrays during recording time |
| Number of network bursts (NB) | Number of network bursts detected in individual arrays during recording time |
| Mean duration of NB | Mean duration of NB calculated as the time from the first spikes to the last spike in individual arrays. |
| CV duration of NB | Coefficient of variance of duration of NB |
| Mean number of spikes per NB | The average number of within-NB spikes in individual arrays during recording time |
| CV number of spikes per NB | Coefficient of variance of number of within-NB spikes in individual arrays during recording time |
| Mean number of contributing channels | Mean number of participating channels per NB in individual arrays during recording time |
| CV number of contributing channels | Coefficient of variance of number of participating channels per NB in individual arrays during recording time |

| | |
|---|---|
| Mean STTC | Mean values across STTC Matrix |
| CV STTC | Coefficient of variance across STTC Matrix |
| Skewness in STTC | Skewness in STTC values from STTC Matrix |

**Table S 3** Normality tests performed to test normality in the data sets of MEA-derived parameters; Four tests were conducted to lower the risk to fail in identifying the proper distribution. Here: 'no' indicates non-normal

| **Data set** | **Statistical Test** | | | |
|---|---|---|---|---|
| | Anderson-Darling | D'agistion - Pearson | Shapiro-Wilk | Kolmogorov-Smirnov |
| wMFR | no | yes | no | no |
| interspike interval | no | no | no | no |
| wMBR | no | no | no | yes |
| Burst duration | no | no | no | no |
| No. of network bursts (NB) | no | no | no | no |
| Mean duration of NB | no | no | no | no |
| Mean no. of spikes p. NB | no | no | no | no |
| Mean STTC | yes | yes | yes | yes |
| Skewness in STTC | yes | yes | yes | yes |

**Table S 4** Identification of pathologically affected networks by test stimulus (STIM). Following steps were performed to select the valid wells by calculation of the relation between the evoked activity upon STIM2 and STIM1 and to check for outliers; According to web-page: Bhandari, P. (2022, November 11). How to Find Outliers | 4 Ways with Examples & Explanation. Scribbr. https://www.scribbr.com/statistics/outliers/ .

1. Calculate the relation between firing activity upon STIM2 and upon STIM1 for each well
   relSTIM = $\frac{evMFR_{norm}^{STIM2}}{evMFR_{norm}^{STIM1}}$
2. Calculate quartile 1 (Q1), quartile 3(Q3) and interquartile range (IQR) of relSTIM across all wells
3. Calculate upper and lower threshold for relSTIM and discard outliers

$$S^{upper} = Q3 + 1.5 \cdot IQR$$

$$S^{lower} = Q1 - 1.5 \cdot IQR$$

#### Supplementary Figures

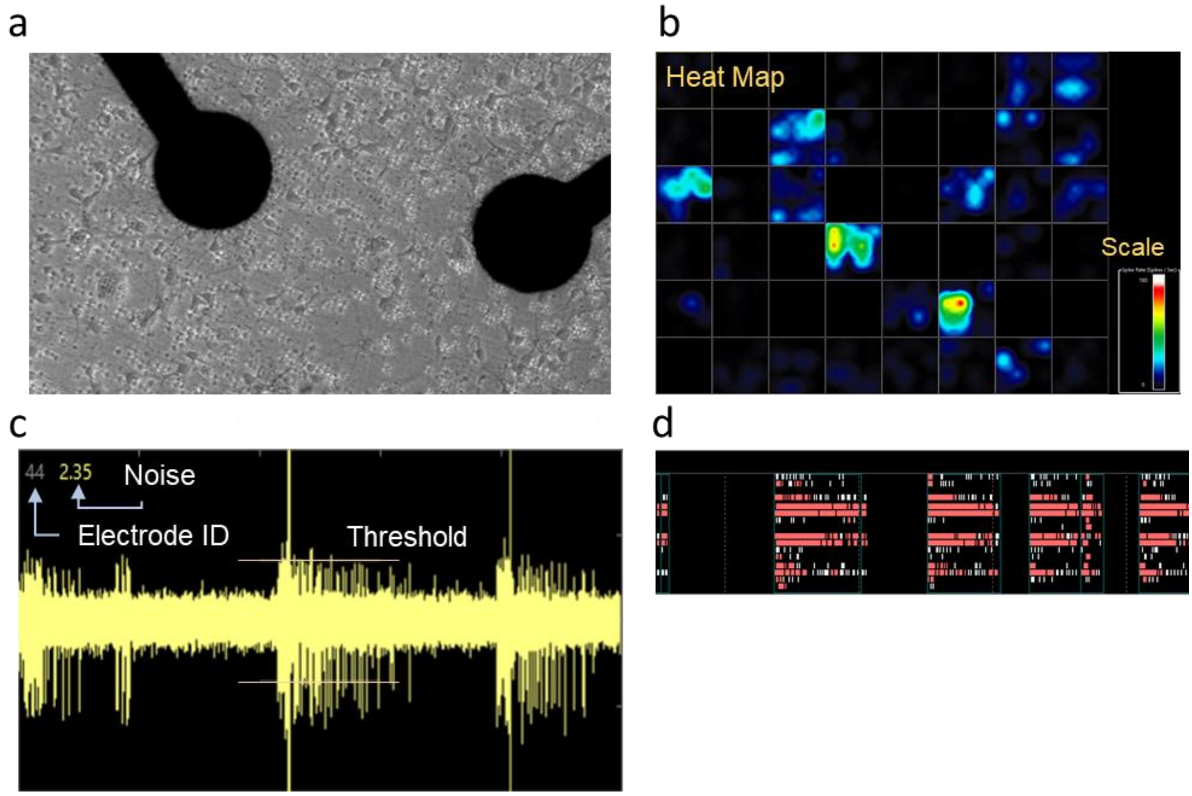


**Fig. S 1** Visualization of cortical cultures derived from *Ptpn11*$^{D61Y}$ mice and activity recordings in Axion Integrated Studio (AxIS) 2.4.2.

**a** Electrodes with neuronal culture on DIV 7 (scale bar: 400 µm); **b** Heat map visualizing network activity across the entire plate on DIV 21, one square corresponds to one well; **c** Continuous Waveform Plots displaying the continuous voltage recording for each electrode on DIV 21. Electrode ID and related noise level in µV are plotted. **d** Raster Plot indicating spikes on DIV 21; spikes within bursts are indicated as red strikes, network bursts are indicated as green rectangles.

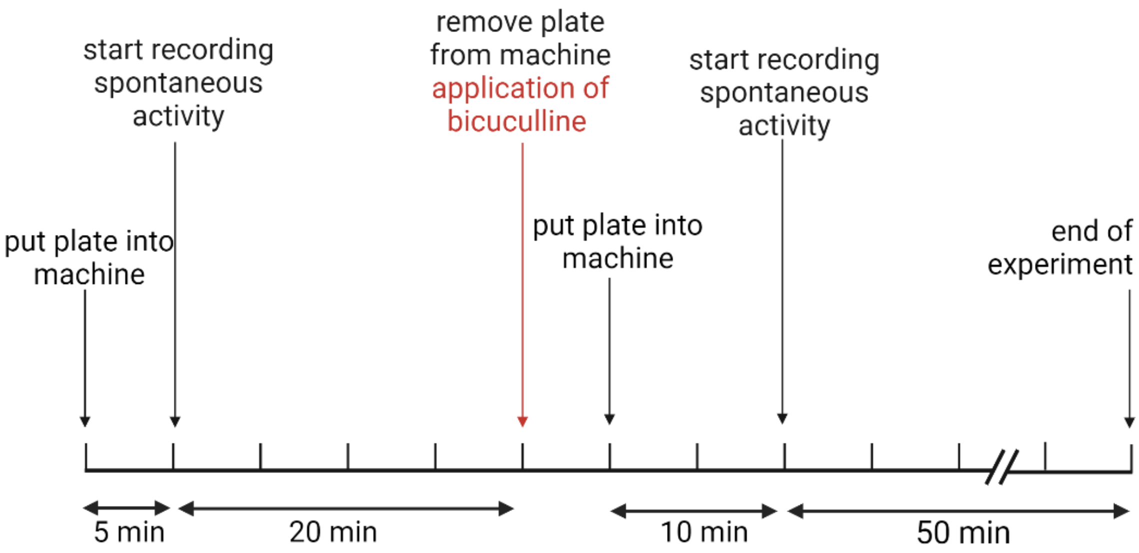


**Fig. S 2** Experimental schema for testing the effect of disinhibition on neuronal activity in *Ptpn11*$^{D61Y}$. Individual steps of experiments are depicted on a timeline. Created with BioRender.com

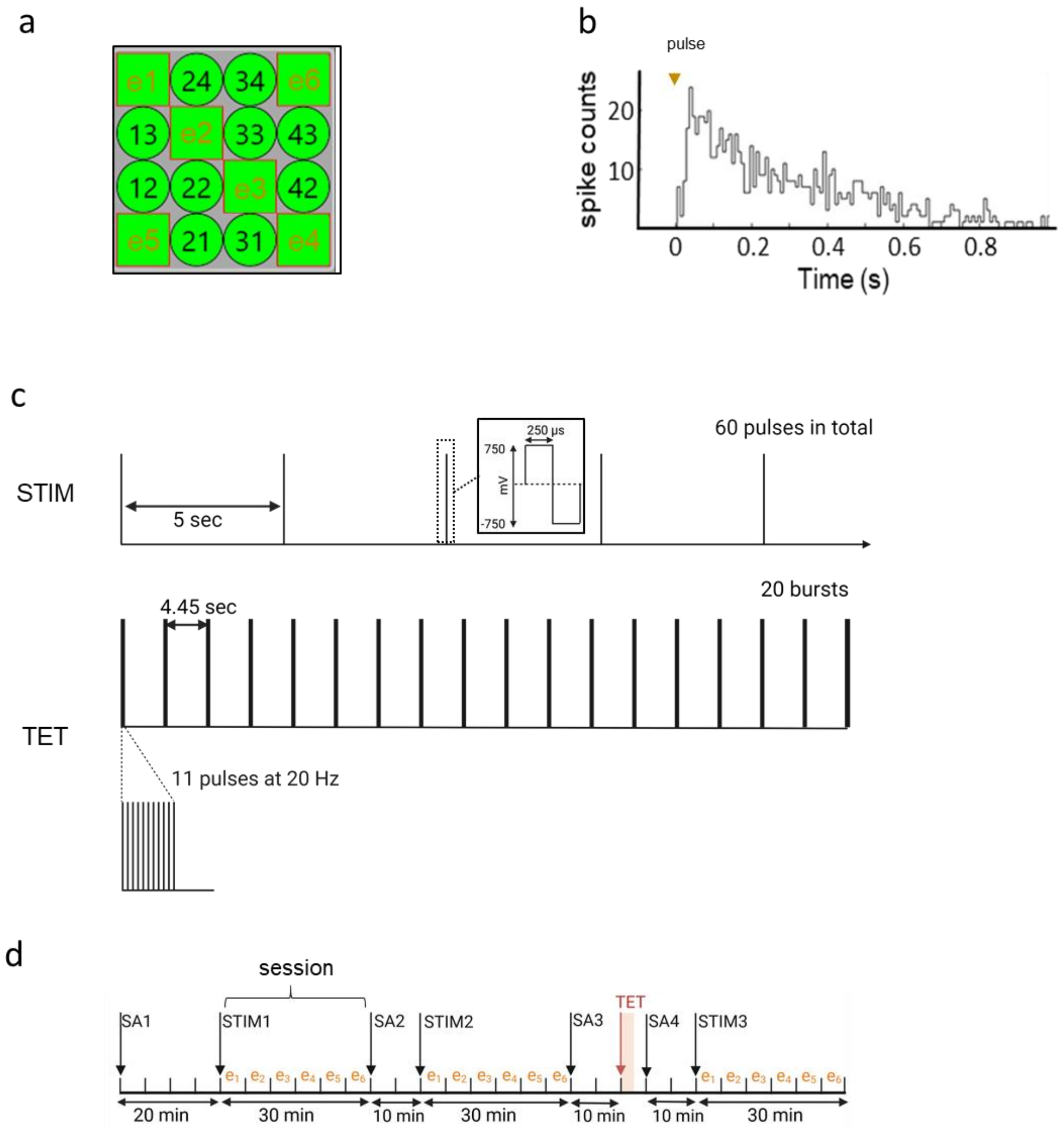


**Fig. S 3: Experimental setting for recording of evoked activity in neuronal networks on MEA**

**a** Schematic illustration of a multi-electrode array (MEA) indicating stimulation electrodes (e1-e6). The electrodes delivered test stimuli (STIM) subsequently (session 1-6) to record evoked activity. **b** Histogram shows number of spikes recorded upon electrical pulse with bin width of 8 ms and total time window of 1 s to sum up spike counts. **c** Stimulation protocols for test stimulus (STIM): 60 pulses at 0.2 Hz were applied to each of the six electrodes (resulting 5 min per electrode, 30 min in total) and for tetanus (TET): 20 bursts consisting of 11 pulses at 20ttzcc Hz were applied to one electrode; all pulses were delivered as biphasic pulse with characteristics depicted in the insert in c. **d** Time line of electrical stimulation protocol including recordings of spontaneous activity (SA) prior and after TET. **(c)** and **(d)** created with BioRender.com

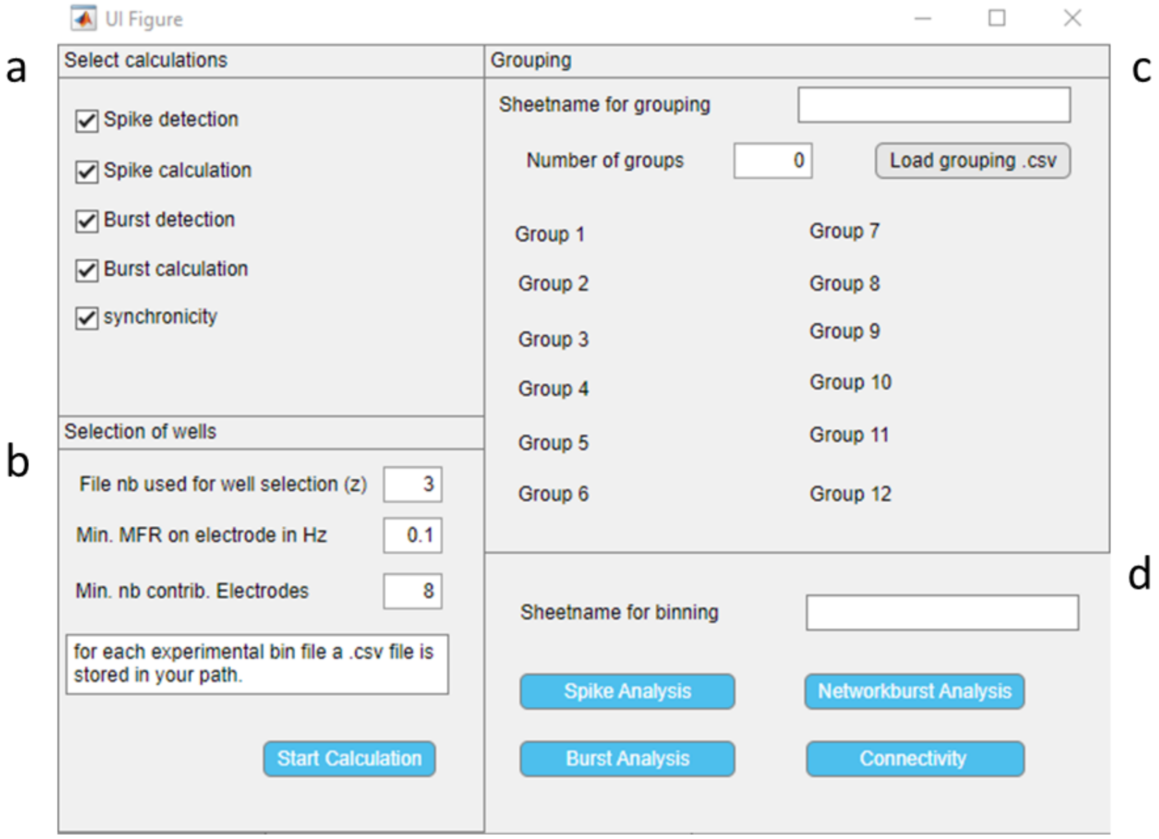


**Fig. S 4** Graphical user interphase (GUI) of custom-written MATLAB based routine to analyse spontaneous network activity in neuronal networks grown on MEA.

**a** Check boxes for selection of operators; **b** Input field for the position (z) in the file list to determine recording/file to select valid wells, input field to determine inclusion criteria for contributing electrodes by minimum MFR (Hz) and for valid wells by minimum number of contributing electrodes per well; **c** Input field for the name of the excel sheet containing grouping information (sheet name for grouping); push bottom to select file and path of grouping file; input field for number of groups; input fields for group names. **d** Input field for name of excel sheet with information about averaging time units; push bottoms to initialize distinct analysis.

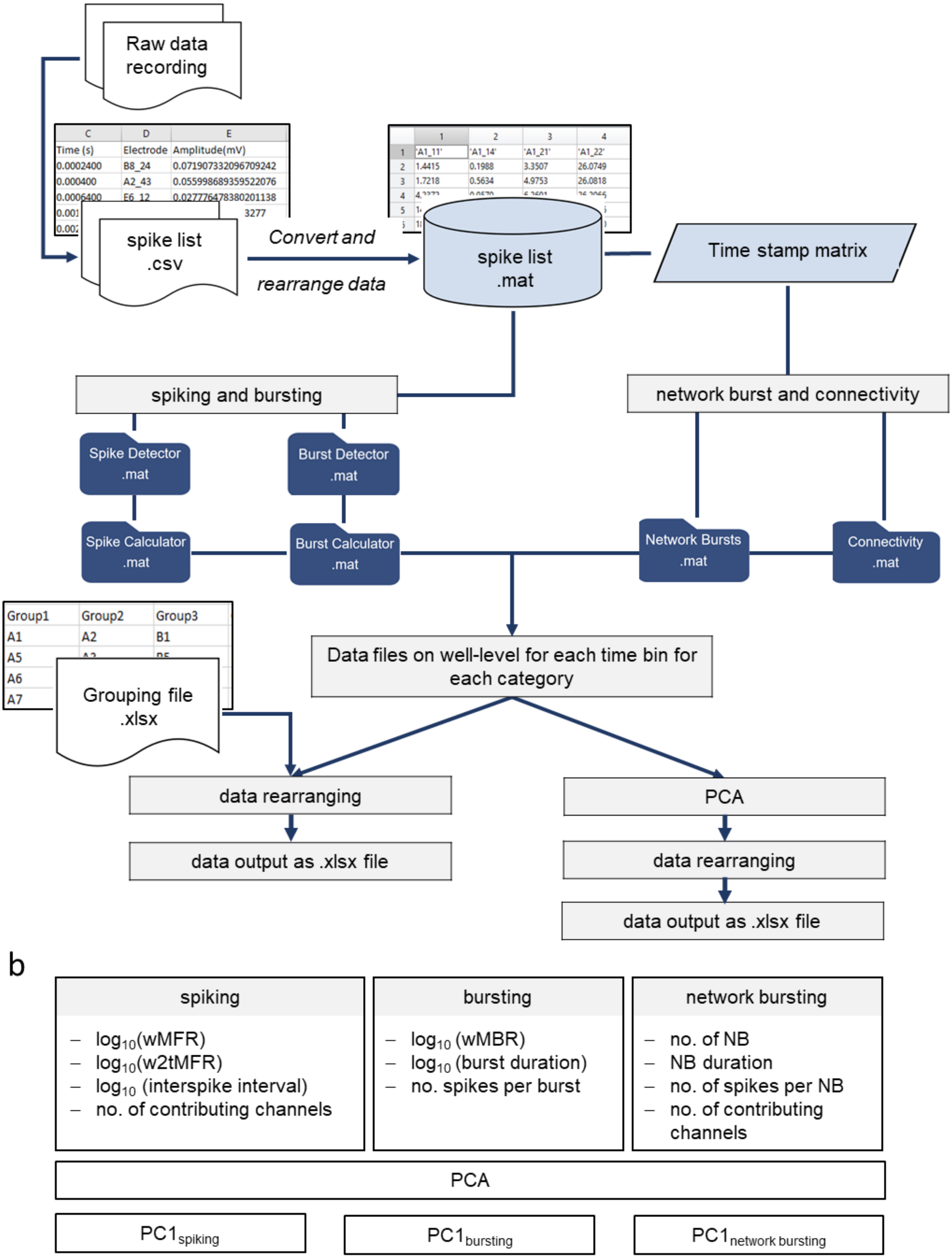


**Fig. S 5** General schemes of data analysis procedures**.**

**a** Automatized analysis of spontaneous activity recordings in time series: raw data is converted to spike list files (.csv) in Axion Integrated Studio (AxIS) 2.4.2. Further processing is performed by custom-written MATLAB software package. Data set is converted and rearranged to a .mat cell array containing the time points of spikes and corresponding electrode name sorted in columns. Parameters describing spiking and bursting were calculated and output on well-level. For network bursting and connectivity parameters, time stamp matrix is calculated beforehand. Then, the parameters are either directly rearranged according to genotype, animal or treatment and averaged according to time units or principle component analysis (PCA) is switched in between. **b**

The functional network features spiking, bursting and network bursting resulting from PCA dimension reduction projected on principle component (PC) 1.

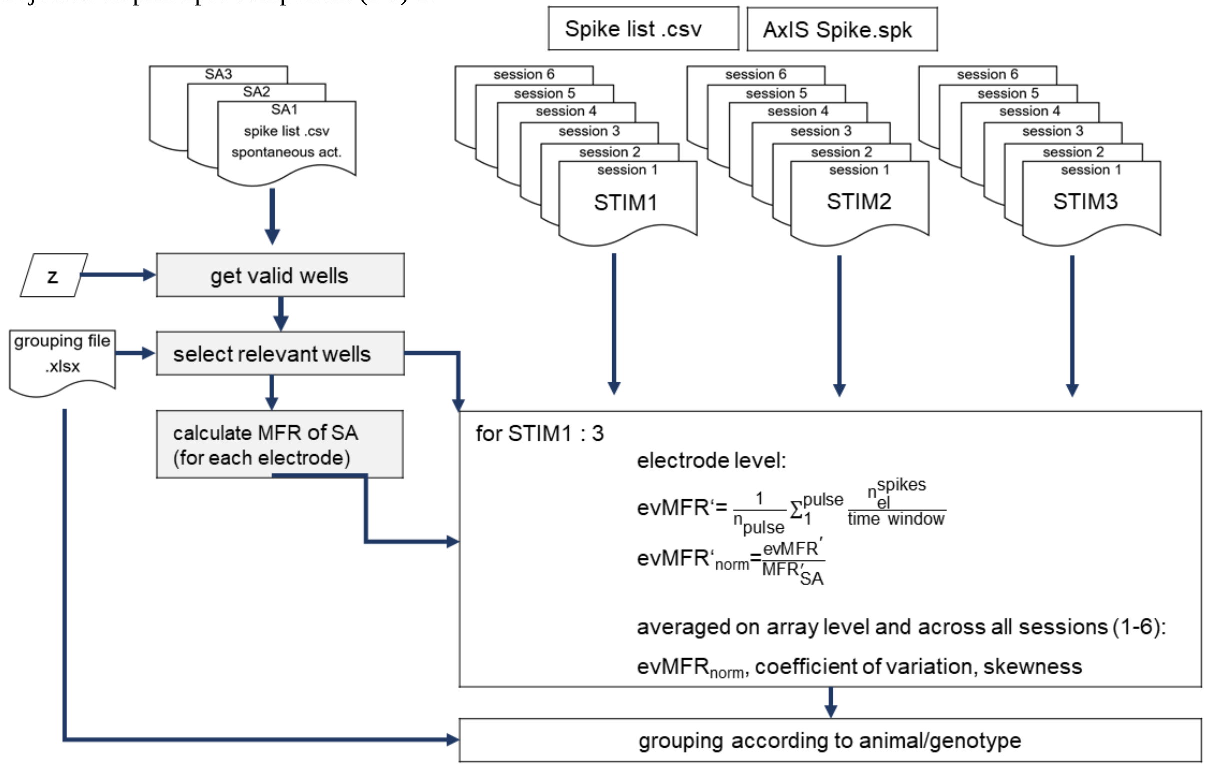


**Fig. S 6** General scheme of automatized analysis of evoked activity by electrical stimulation.

Input data derives from Axion Integrated Studio (AxIS) 2.4.2. Input consists of .csv spike list files, containing information about electrode name and time point of each spike chronologically. Spike list files are entered from spontaneous activity recordings and evoked activity recordings, as well. Information about electrical stimulations is given in AxIS Spike .spk files. Data processing is performed by custom-written MATLAB functions. A session comprises triggering of six electrodes in a row with test pulses at 0.2 Hz for 5 min each. Selection of valid wells performs on a spontaneous activity (SA) recording prior to test stimulus (STIM) 1 defined by z (number of the related file in the folder containing all recordings) inserted by the user. Groups of wells are defined in the .xlsx grouping file. MFR from spontaneous activity recordings is used to normalize the evoked activity on each electrode.

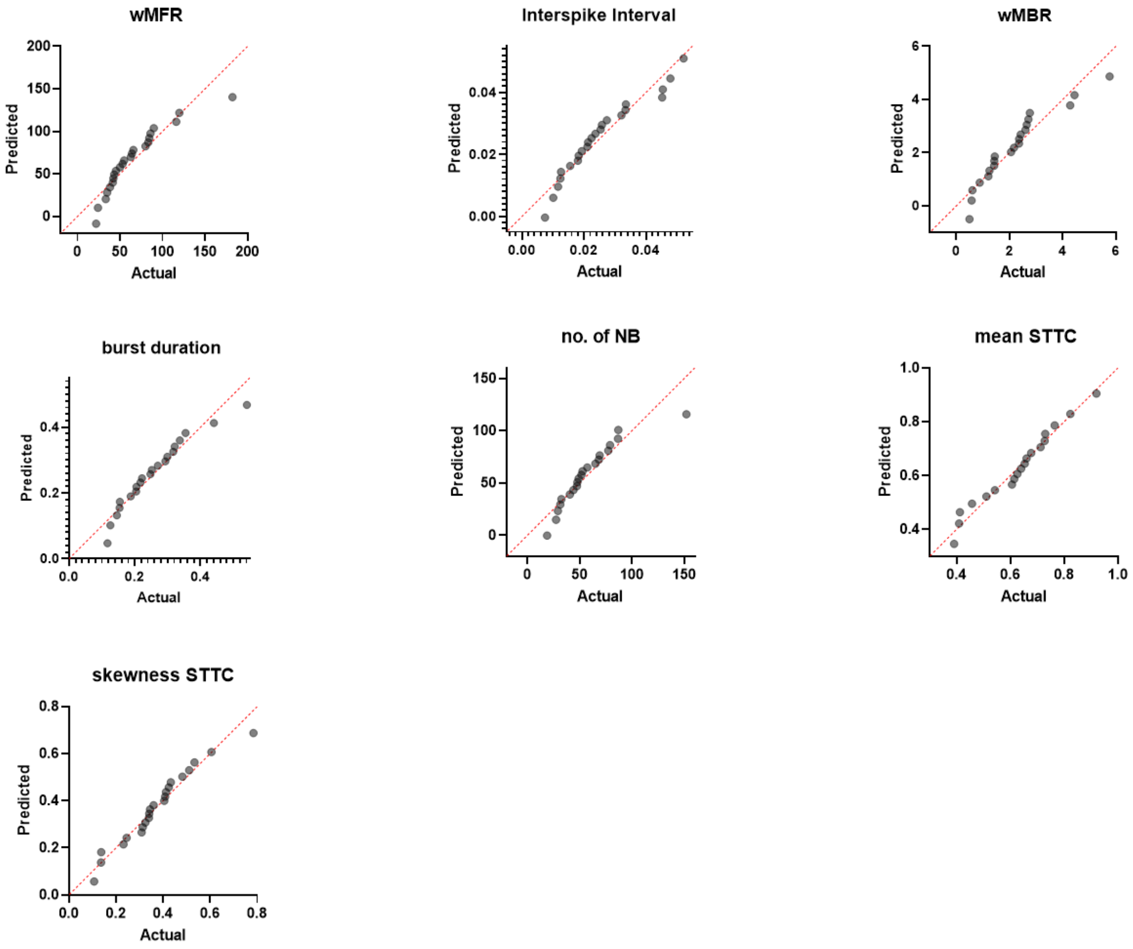


**Fig. S 7** Distribution of data sets as normal QQ plots referred to distinct parameters from time series experiments derived from wild type and knock in of *Ptpn11*$^{D61Y}$.

QQ plots showing predicted residual versus actual residual. A data point represents the averaged value across the wells belonging to one preparation/animal on one plate (preparation level). Calculation is based on data recorded on DIV21;

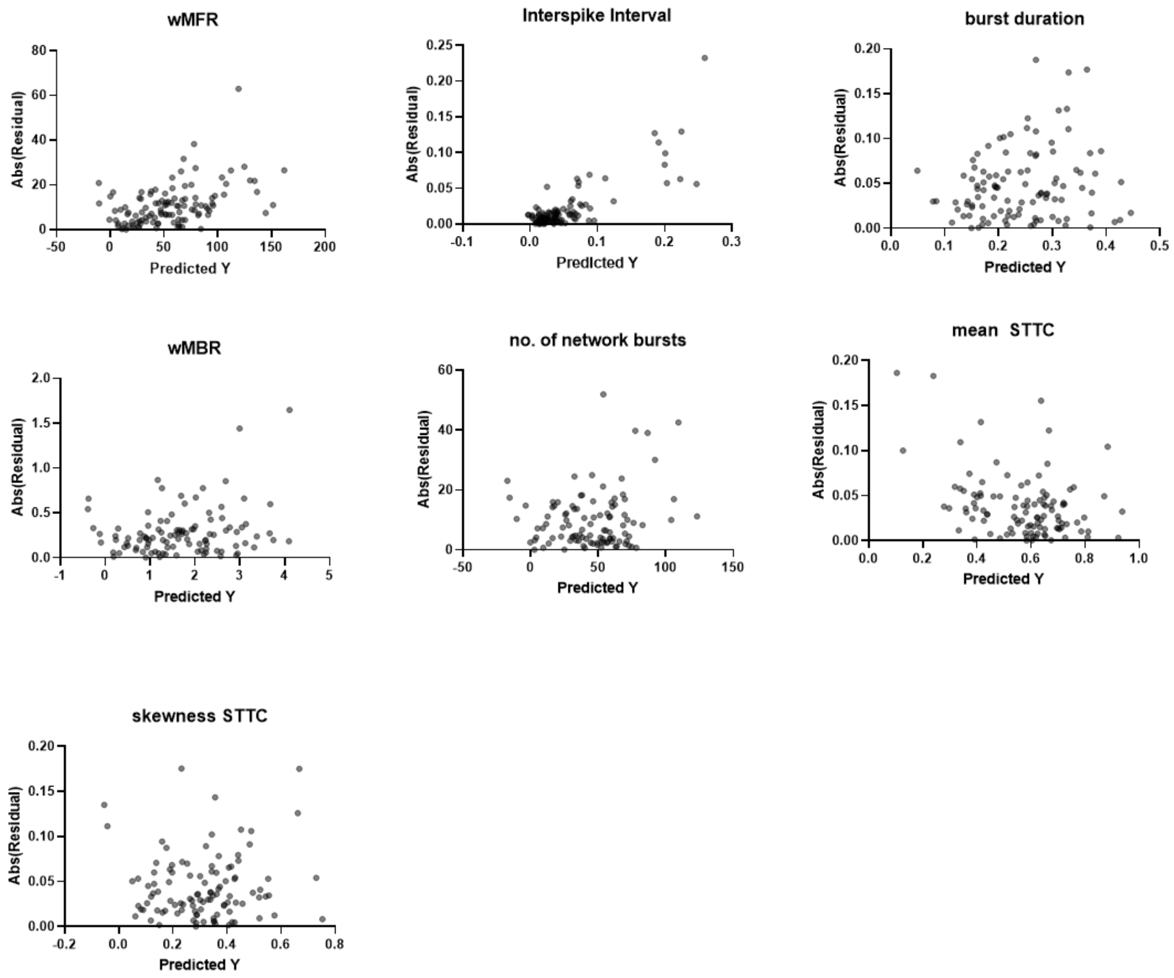


**Fig. S 8** Homoscedasticity plots for recorded individual features describing spontaneous activity in longitudinal recordings in *Ptpn11*$^{D61Y}$.

Absolute values of residual versus predicted values of several parameter describing spontaneous network activity. A data point represents the averaged value across the wells belonging to one preparation/animal on one plate (preparation level). Calculation is based on the data recorded on DIV21;

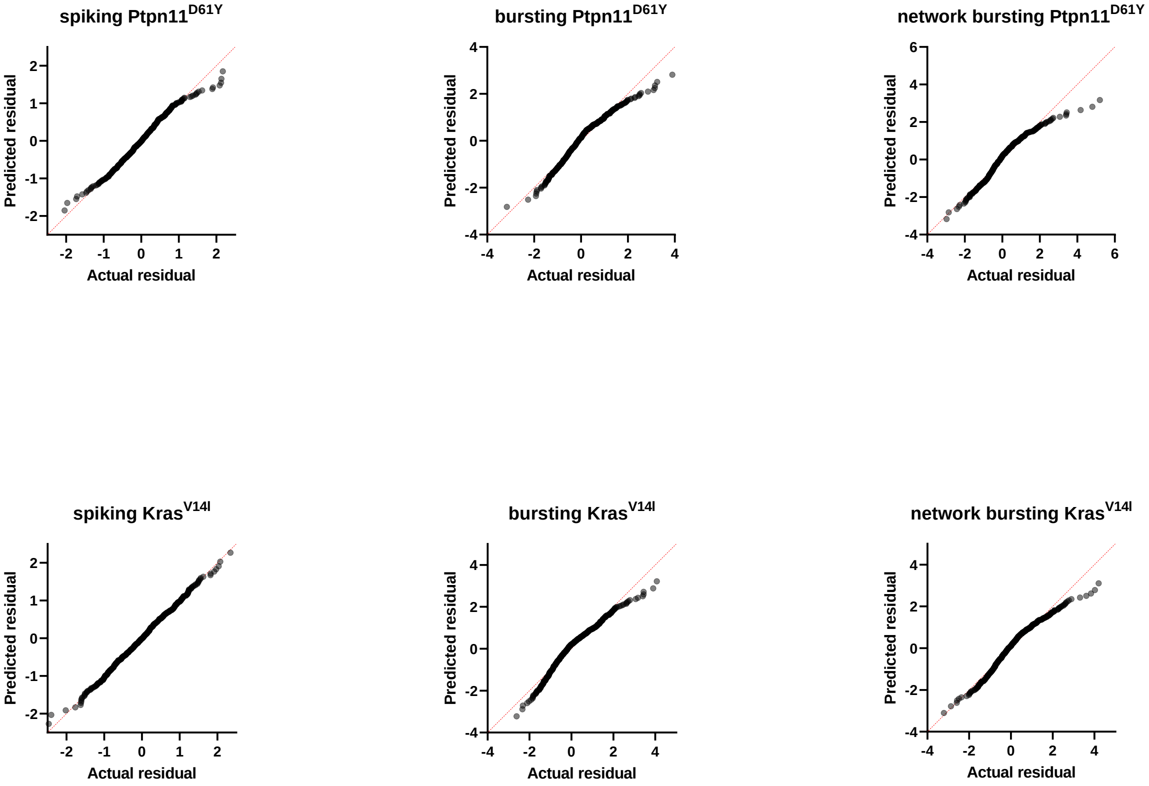


**Fig. S 9** Distribution of PC1 projected data sets as normal QQ plots for *Ptpn11*$^{\mathrm{D61Y}}$ and *Kras*$^{\mathrm{V14I}}$. Projection of feature vectors describing the fields spiking, bursting and network bursting onto principle component (PC) 1. A data point represents one well (well-level).

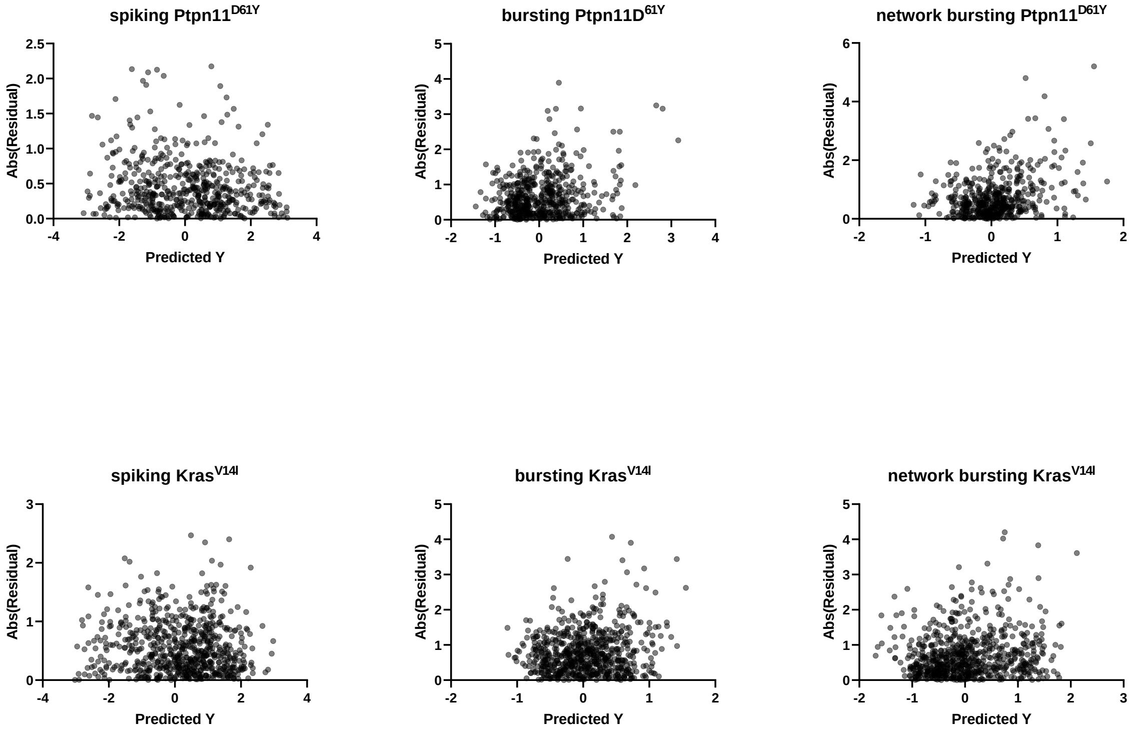


**Fig. S 10** Homoscedasticity plots for recorded PC1 projected data sets for *Ptpn11*$^{D61Y}$ and *Kras*$^{V14I}$.
Absolute value of residual versus predicted values of the projection of feature vectors describing the fields spiking, bursting and network bursting onto principle component (PC) 1. A data point represents one well (well-level).

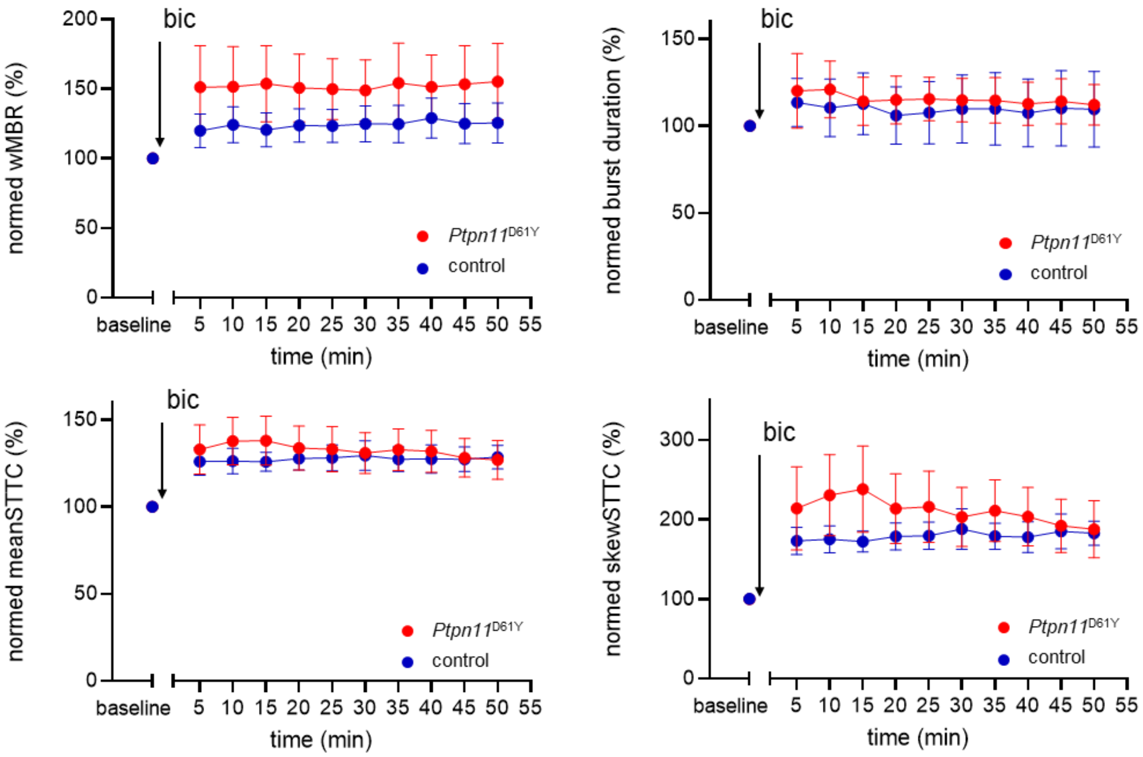


**Fig. S 11** Effect of disinhibition on neuronal activity in *Ptpn11*$^{D61Y}$ with time.

Line graphs with point representing mean ± SEM of several parameters describing network activity as a function of time prior and after application of bicuculline (bic) on DIV 33. Baseline values (averaged values over a time interval of 20 min prior bic treatment) were set as 100 %, values upon bic treatment were related to baseline.

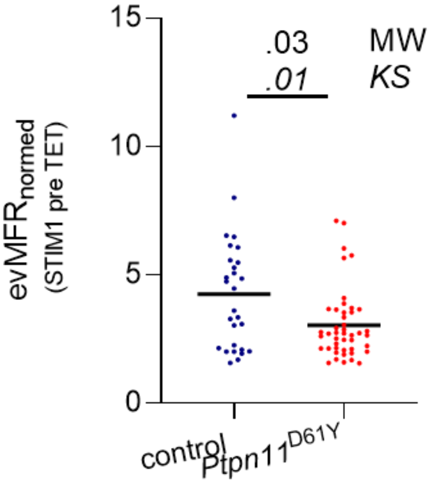


**Fig. S 12** Assessment of evoked activity in *Ptpn11*$^{D61Y}$ RASopathy model. Scatter dot plot demonstrating. evMFRnormed upon stimulation pre TET (STIM1), black lines indicate median values, one data point refers to one well. Significance was tested by Mann-Whitney (MW) to check differences in mean values and Kolmogorov-Smirnov (KS) to check differences in data distributions.

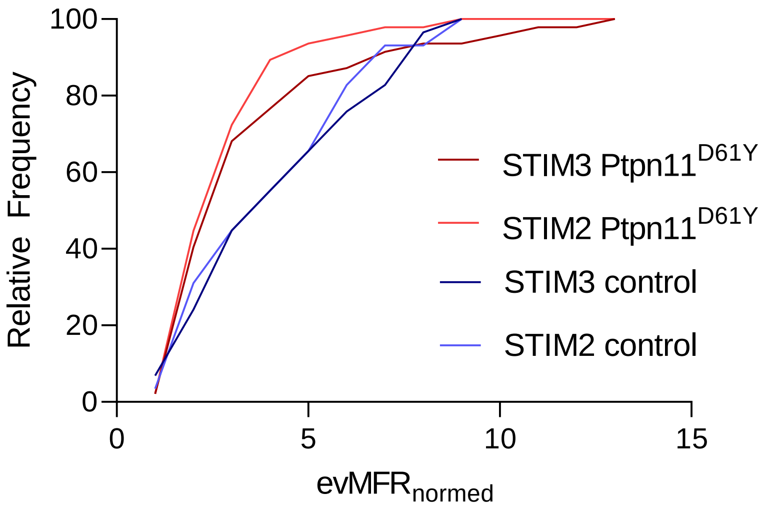


**Fig. S 13** Relative frequency for $evMFR_{normed}$ upon STIM2 and STIM 3 for control and *Ptpn11*$^{D61Y}$
